# Over-the-horizon extinction risk assessment reveals rapidly shifting geographic and taxonomic priorities for conservation

**DOI:** 10.64898/2026.04.13.718320

**Authors:** Alex Slavenko, Marcel Cardillo, Lindell Bromham, Xia Hua, Ben C. Scheele

## Abstract

A central challenge in conservation is understanding how climate change interacts with other global-change drivers to shape future species extinction risk, threatened-species hotspots, and the effectiveness of protected areas. Here, we use an integrated over-the-horizon forecasting framework to jointly model changing species’ range dynamics and shifts in extinction risk for 1,914 Australian terrestrial vertebrates to 2100. Our approach links ensemble species distribution models with machine-learning–based automated threat assessment, incorporating species traits, changing distributions of invasive species, and projections of land use and human population density. Under a high-emissions scenario, up to 109 species are projected to lose all climatically accessible habitat by 2100 and the number of threatened species is predicted to increase, while under a moderate emissions scenario (SSP1.26) the number of threatened species remains relatively stable, and up to 19 lose all climatically accessible habitat. Spatially, threatened-species richness becomes increasingly concentrated in southeastern Australia. These shifts elevate the representation of threatened species within existing protected areas, largely because extinctions and range contractions occur disproportionately outside protected areas. Our results highlight that the identity of at-risk species and the occurrence of threatened-species hotspots will change dramatically, underscoring the need for forward-looking conservation strategies that anticipate future biodiversity patterns.

## INTRODUCTION

Biodiversity loss in the Anthropocene is driven by multiple interacting threats^1,2^. Yet most forecasts of future extinction risk focus on climate-driven shifts in species distributions. As well as climate, extinction risk also changes in response to changes in land use, human population density, and other threatening processes^3,4^, and the effects of these processes are mediated by species biological traits^5^. Although recent developments have advanced the modelling of species responses to climate change, substantial uncertainty remains about how climate-driven range shifts interact with other global-change drivers and species’ intrinsic biological traits to influence future threat status^5,6^. Addressing this gap requires an integrated understanding of how multiple threats and risk factors interact to determine how species’ geographic ranges and extinction risk are likely to change through time.

One of the major challenges created by changing patterns of extinction risk is that conservation priorities—both species-level and spatial—are unlikely to remain stable. For example, protected areas (PAs) are a central pillar of conservation, with locations with high concentrations of threatened or endemic species prioritised for the establishment of new PAs^7,8^. However, given climate-driven distributional change and shifting threats, the identity of threatened species and the location of future biodiversity hotspots may change substantially through time^9^. Past studies have reached mixed conclusions about whether PA networks will lose or gain effectiveness under climate change. Earlier research concluded that PAs would decline in effectiveness because they lose more species than they gain under climate-driven range shifts^9,10^. However, more recent work has reported that PA effectiveness has recently increased^11–13^ or is predicted to increase under modelled future scenarios^14–18^. These mixed findings highlight that conservation decisions based on present-day patterns risk being misaligned with future biodiversity needs—a challenge that can only be addressed by forecasting the joint dynamics of species distributions, threatening processes, and extinction risk.

Here we present an integrated approach to forecasting future extinction risk that combines ensemble species distribution modelling with information on species biology and projected changes in threatening processes (Figure⍰1). Our approach uses a machine-learning–based automated assessment framework to classify species by their predicted IUCN Red List status to the end of this century. This automated assessment method was developed to predict the threat status of unclassified reptile species^6^ but has not previously been applied to forecasting future changes in threat status. We use our integrated approach to investigate changes in the distribution and extinction risk in terrestrial Australian vertebrates, a globally unique and diverse continental assemblage of >2,200 species. We quantify temporal changes in the number and proportion of threatened species across the four vertebrate classes and assess how spatial patterns of threatened-species richness are expected to shift. We then evaluate how effectively the current PA network is predicted to represent individual threatened species and threatened-richness hotspots under future scenarios, considering climate-driven range shifts and projected changes in species threat status simultaneously. Finally, we analyse changes in cross-taxon congruence in hotspot–PA overlap to evaluate whether future priority areas are likely to become more spatially concentrated or dispersed.

**Figure 1.**
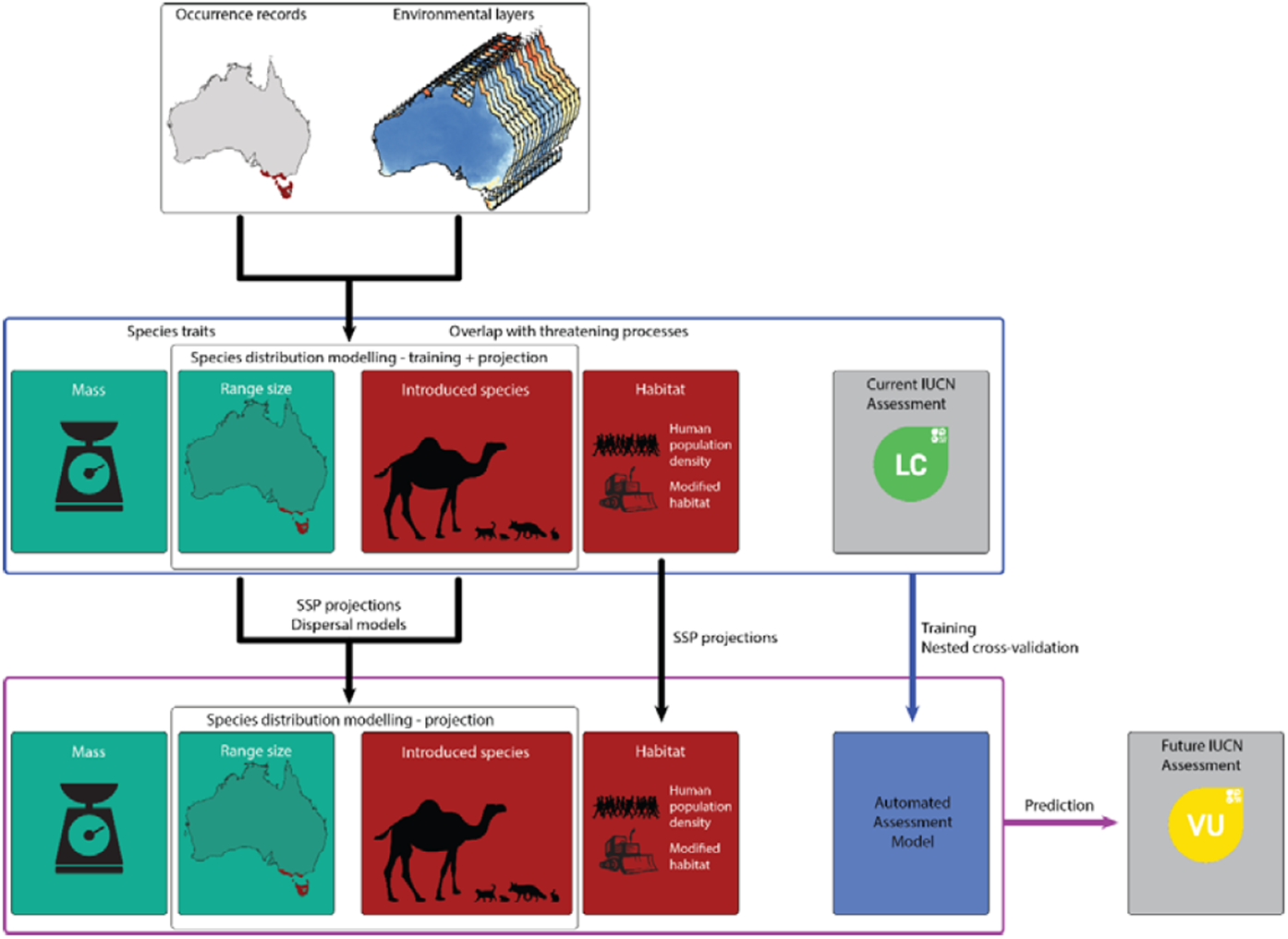
Workflow for our over-the-horizon framework to predict extinction risk. Curated occurrence records are coupled with environmental layers (bioclimate, soil, etc.) to train species distribution models (SDMs), both for assessed native species and invasive species. These are used to generate features for machine learning-based automated assessment models, consisting of species traits (mass and range size, the latter projected from the fitted SDMs) and overlap with threatening processes (SDM projections of introduced species ranges and sourced layers of human-population density and modified habitat). These features, coupled with current IUCN threat assessments, are used to train an automated assessment model using nested cross-validation. Then, future projections of climate layers under different Socioeconomic Pathways are used to project the SDMs and generate future range maps for the native species and invasive species under different dispersal scenarios. Future projections of range size and overlap with threatening processes are used to make new predictions of future IUCN threat assessments using the automated assessment model.

## RESULTS

### Predicted changes in species threat status: taxonomic patterns

We implemented a novel, integrated workflow for forecasting future changes in species threat status, by combining independently projected changes in geographic range size of native and invasive species (as a threatening process) from SDMs based on 5.6 million occurrence records, with projected changes in human population density and land use, and biological and geographic predictors of extinction risk. We applied an XGBoost automated assessment model^6^ on this integrated dataset for 1,914 Australian terrestrial vertebrate species, achieving high classification accuracy (0.914) when distinguishing threatened (VU, EN/CR) versus non-threatened species (NT, LC) and Near Threatened vs. Least Concern species (0.937), but lower accuracy (0.604) when distinguishing between the Threatened categories (Table S1). The most important predictors of current threat status were geographic range size and body mass, followed by habitat-related features and overlap with invasive species (Figure S1, Tables S2–S3).

Under the more pessimistic SSP5.85 emissions scenario, the number of species in each of the four vertebrate classes (mammals, birds, reptiles, amphibians) that are classified in a Threatened (VU, EN/CR) category is predicted to increase throughout the century (Figure 2A, Table S4). This is also the case under the optimistic SSP1.26 scenario when no dispersal is allowed, but with limited or limitless dispersal, the number of threatened species is predicted to remain relatively stable. Our models suggest that stepwise progression through the Red List levels is common: the higher a species’ current threatened level is, the more likely it is to be threatened (or go extinct) in the subsequent timestep (Figure 2B, Table S5). However, the changing taxonomic patterns of extinction risk are also driven by species moving through the levels of the Red List in various directions, *i*.*e*., species moving from unthreatened to threatened, threatened to unthreatened, and from threatened or unthreatened to extinct (Figure S2). Changes in range size led to species predicted to become less threatened, on average (Figure S3). However, changes in most other predictors tended to cause species to become more threatened, particularly those predictors with high importance (Figure S1). Patterns differed between taxa: for example, overlap with rabbits was a stronger driver of higher threat predictions for reptiles than other classes, whereas overlap with cats was a stronger driver of higher threat predictions for amphibians than other classes (Figure S3).

**Figure 2.**
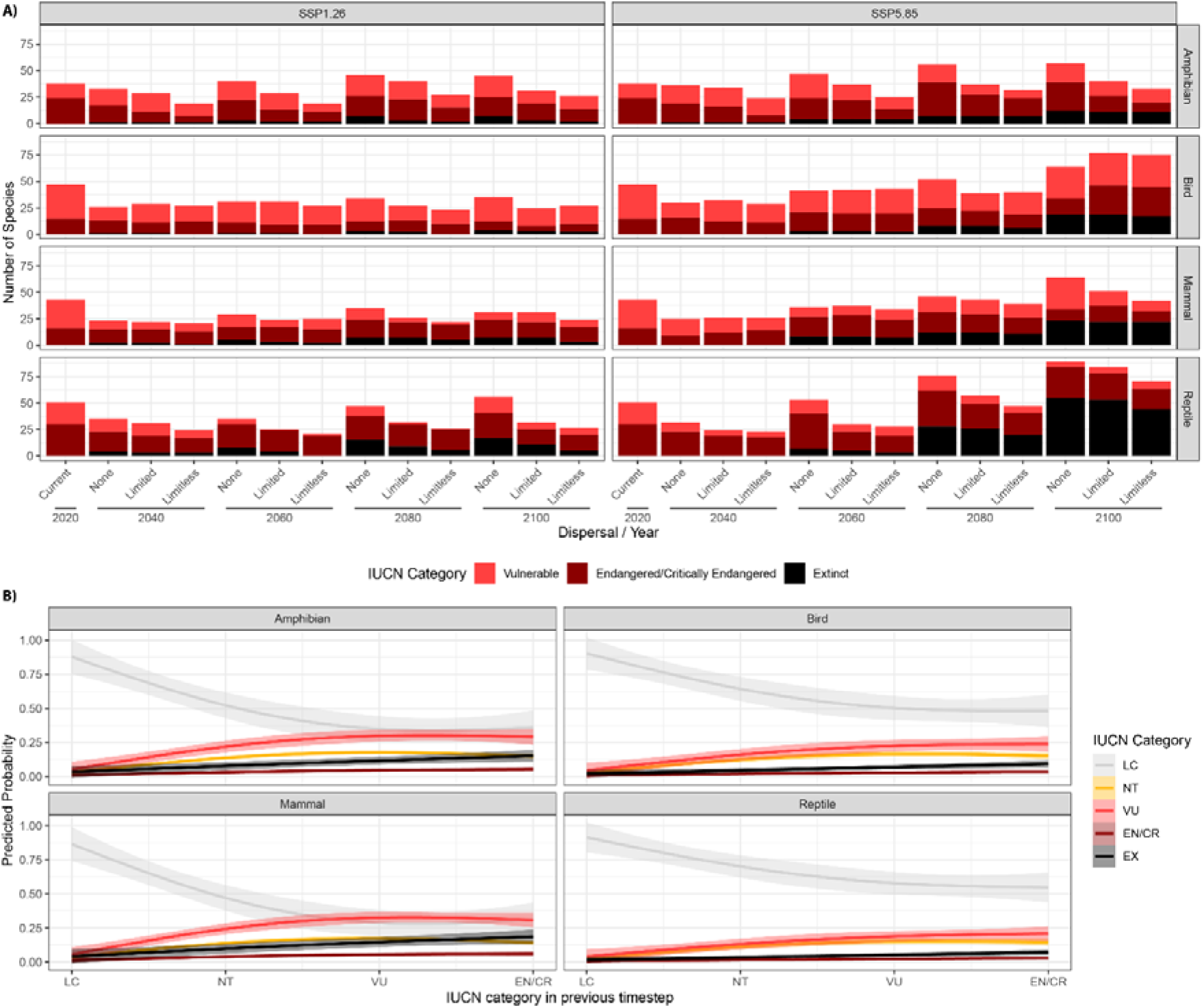
Future threat levels of Australian terrestrial vertebrates. A) Predicted numbers of species in different threatened categories (Vulnerable, Endangered/Critically Endangered) under two different Shared Socioeconomic Pathways (SSP1.26, SSP5.85) and three dispersal scenarios (None, Limited, Limitless) in 20-year timesteps from 2020-2100. Extinct species are coloured black and include species that went extinct in previous timesteps. B) Predicted probability of assignment to each IUCN threat category as a function of the threat category in the previous timestep. Mean predicted probabilities and 95% CI are estimated from predictions across all Shared Socioeconomic Pathways and dispersal scenarios.

The models also predict an increase in the number of extinctions (Figure 2A), which are inferred when a species’ modelled climatic suitability from one time period to the next results in no climatically suitable accessible areas. Under SSP1.26, our models predict 2–7 species of amphibians, 2–4 species of birds, 6–7 species of mammals and 9–17 species of reptiles could go extinct by the end of the century. Under SSP5.85 all groups are predicted to experience a sharp increase in extinctions late in the century, in the period 2080–2100 (Figure 1A), with 11–12 species of amphibians, 17–19 species of birds, 22–23 species of mammals, and 45–55 species of reptiles potentially extinct by the end of the century. Under both scenarios, all but one of these predicted extinctions are species endemic to Australia, in addition to the loss of the shrill whistlefrog (*Austrochaperina gracilipes)* from the Australian mainland (the species is also present in New Guinea, which is not included in our models).

The spatial distribution of predicted extinctions reflects the patterns of range contraction. Although extinctions are predicted in almost every biome by 2100, the biomes of southern and southeastern Australia are expected to carry a disproportionately high share of species losses (Figure 3A, Figure S4). Under SSP5.85, three extinction hotspots emerge (Figure S4): the dry monsoonal Gulf Country of northern Australia, the Victorian and Murray Mallee, and Tasmania, which is predicted lose 2–27 endemic species, depending on the modelled dispersal scenario.

**Figure 3.**
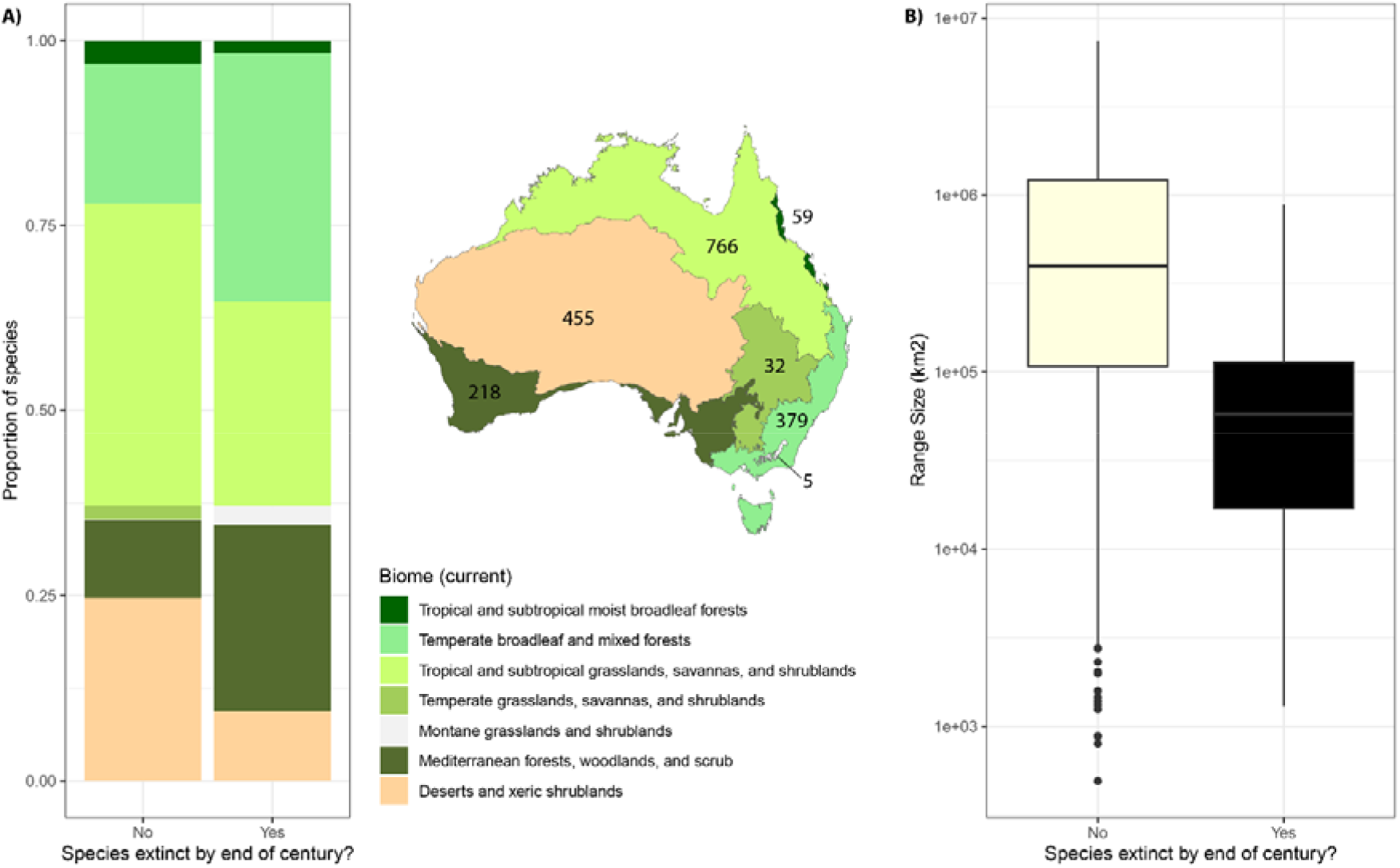
Distribution of Australian terrestrial vertebrate species predicted to go extinct by the end of the century. A) Bar plots representing for species predicted to survive (left) or go extinct (right) the proportion of species currently occurring in each biome. Extinct species are considered across all modelled scenarios of climate change (two SSPs and three dispersal scenarios), thus representing the most pessimistic scenario. Numbers next to each biome in the centre map represent the total number of species with the majority of their extent distribution in each biome. B) Box plot showing the distributions of current range size (in km^2^; log_10_-transformed) of species predicted to survive or go extinct under all modelled scenarios of climate change.

### Shifting species distributions under climate change

Shifts in the centroids of species’ geographic ranges (Figure S5) are driven by leading-edge expansion of the range boundary on one side, trailing-edge contraction of the boundary on another, or both. In northern Australia, expanding areas of climatic suitability to the southeast mean that on average, species ranges are expected to expand (Figure S6). In southern Australia, hard limits to range expansion at the edges of the continent mean that on average, species ranges are expected to contract (Figure S6).

The predicted range changes differ between taxa and vary over time. The pattern of northern expansion of ranges is strongest earlier in the century (2020–2040), while the pattern of southern range contraction becomes stronger as the century progresses (Figure S6). Range changes are relatively evenly split between contractions and expansions in the first half of the 21^st^ century, but contractions become more prevalent and of greater magnitude following 2060 (Figure S7).

### Changing spatial patterns of threatened species richness

Current threat hotspots (top 10^th^ percentile of threatened species richness) for the four classes are located primarily in the regions of highest total species richness along Australia’s eastern seaboard, with large hotspots for mammals and reptiles also found in northern Australia, and for mammals in southwestern Australia (Figure S8A). Under all emissions and dispersal scenarios these hotspots are predicted to shift towards the south, to become concentrated in the southeastern extremity of the Australian mainland (amphibians, mammals & reptiles), and the eastern/southeastern coastal strip (birds), by the end of the century (Figure S8B–D). This general shift pattern across the four classes is reflected in the hotspots for combinations of one, two, three and four classes, which are predicted to contract from northern Australia and the central-east coast region towards the southeastern extremity of Australia, particularly under SSP5.85 (Figure 4A).

**Figure 4.**
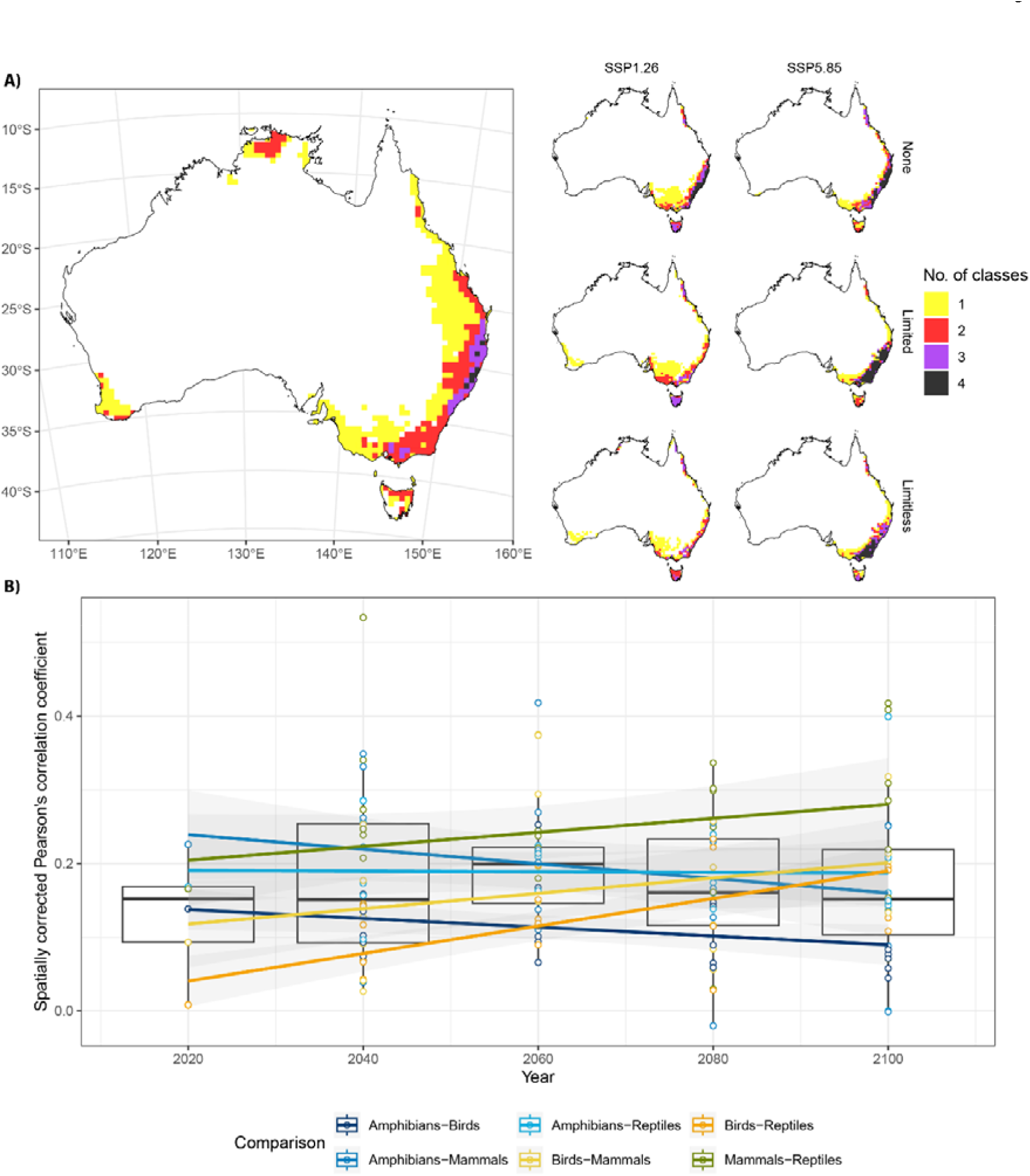
Changes in congruence of threat hotspots. A) Maps representing hotspots of threatened species richness for the four classes of terrestrial vertebrates. Colours represent degree of congruence, *i*.*e*., how many classes have hotspots in that cell. Left panel shows current hotspots based on automated assessment of 1,914 species, and right panels show projected hotspots at 2100 under two different Shared Socioeconomic Pathways (columns) and three different dispersal scenarios (rows). B) Box plots showing the degree of congruence in threatened species richness under two different Shared Socioeconomic Pathways (using the Limited dispersal scenario) over 20-year timesteps (2020 representing present conditions), calculated using spatially corrected Pearson’s correlation coefficients. Coloured lines represent trends in different pairs of taxa (*e*.*g*., the orange line shows increasing congruence between birds and reptiles).

Some more idiosyncratic patterns of hotspot shifts are also evident. In birds and mammals, the concentration of threatened species on the island of Tasmania intensifies to hotspot level under SSP1.26 (Figure S8B–D). This hotspot intensification is not seen under SSP5.85, however, due to a predicted large wave of extinctions of endemic Tasmanian threatened species leaving mostly non-threatened species remaining. Hotspot areas for mammals and reptiles in northern Australia are expected to contract and disappear as species expand their distributions southwards and reduce their threat status (Figure S8B-D). In amphibians and, to a limited extent, in mammals, the Wet Tropics in the far north-east of Australia emerge as a threatened species hotspot (Figure S8B–D). Because classes are predicted to respond differently to climate change, these changes lead to an increase in spatial congruence of threatened richness hotspots of mammals and reptiles with birds, and a reduction in spatial congruence between amphibians and mammals (Figure 4B; Table S6).

The area of threatened species richness hotspots, as a percentage of Australia’s land area, is predicted to decrease by 2100 for single-class hotspots (from 9.13% to 3.43–7.42%, summed across all four classes and depending on SSP and dispersal scenario), and for hotspots shared by two classes (4.42% to 1.24–2.94%) (Figure 4A). However, as threatened species richness in general becomes more concentrated in southeastern Australia, hotspots shared by three classes (1.06% to 0.67– 1.58%) and all four classes (0.21% to 0–2.95%) are predicted to increase in size under most scenarios.

### Overlap between threatened species and Protected Areas

Threatened species richness hotspots are predicted to contract and become concentrated in southeastern Australia, especially mainland upland areas and the island of Tasmania, areas which have relatively high PA coverage. This leads to a predicted increase in overlap between hotspots and current protected areas by 2040, and throughout the century (Figure 5A). For species predicted to be threatened, we predict increases in the mean overlap of species ranges with PAs, but no increase for predicted non-threatened species (Figure 5B). There are also predicted increases in the proportion of threatened species that overlap substantially with the current PA network, while for non-threatened species the overlap increase is only predicted under a zero-dispersal model (Figure S9). Under zero-dispersal models, the number of species predicted to reach 50% overlap of their ranges with PAs is greater than the number of species whose level of overlap will decrease to below 50% (Figure 6, upper panels). However, because the no-dispersal model does not allow range expansion, this essentially means that range contractions are predicted to be more extensive outside PAs than within them. This pattern is more variable and inconsistent under the models that allow limited or limitless dispersal (Figure 6, middle and lower panels), since under these scenarios ranges can also expand out of PAs, reducing species’ coverage within PAs, or currently un-represented species can expand their ranges into PAs, increasing their coverage. For amphibians and reptiles, more species are expected to lose the 50% level of overlap with PAs than gain them, reducing the overall protection for these two taxa. Conversely, many more bird species are expected to reach the 50% overlap level than lose it.

**Figure 5.**
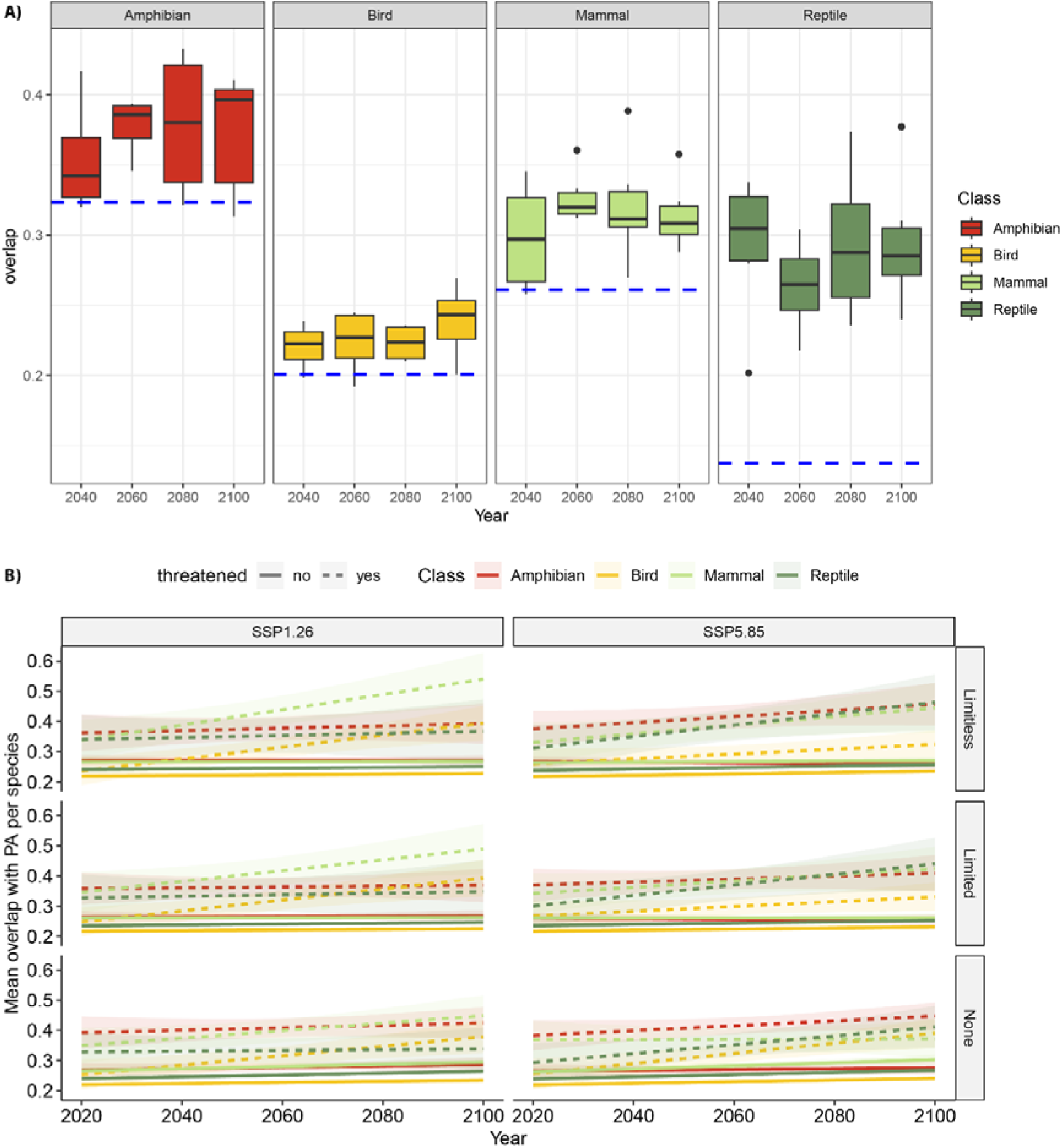
Degree of overlap with protected areas. A) Box plots showing the degree of overlap between threatened species hotspots and protected areas under all considered climate change scenarios over 20-year timesteps, calculated as the proportion of hotspot area overlapping with any protected area. The horizontal blue dashed lines represent current levels of overlap with protected areas based on automated assessment of 1,914 species. B) Mean levels of overlap with PAs predicted through time, faceted by different Socioeconomic Pathways and dispersal scenarios. Solid lines represent predictions for non-threatened species, and dashed lines represent predictions for threatened species.

**Figure 6.**
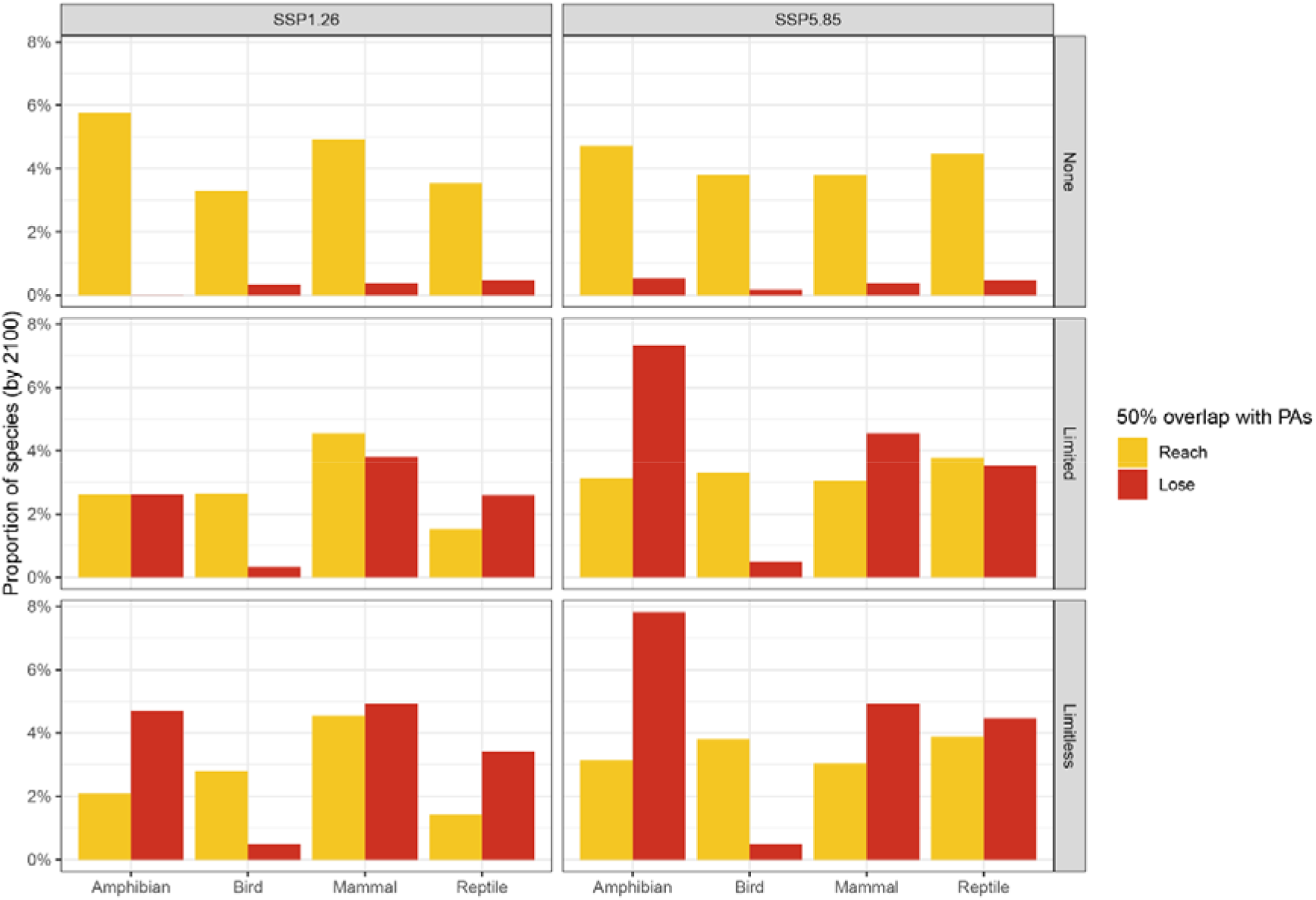
Changes in levels of protection for threatened species. Bar plots showing the proportion of threatened species in each terrestrial vertebrate class that either drop below the 50% overlap with PA level (in red) or gain that protection level (in gold). Results are faceted by different Socioeconomic Pathways and dispersal scenarios.

## DISCUSSION

Methods for forecasting species extinction risk over decadal timescales are becoming increasingly important as conservation shifts to being more proactive^4,6,19,20^. Robust extinction-risk forecasting requires methods that consider the many factors that are likely to influence a species’ changing threat status: this includes not only climate change driven distribution shifts, but also projected changes in threat such as human population, land use, and invasive species, and the way that species’ intrinsic biology mediates responses to threats. We have presented the first (to our knowledge) integrated forecasting methodology that combines these factors to simultaneously examine shifts in species distribution and changes in threat status. Our models predict profound changes in the continent-wide taxonomic and spatial patterns of extinction risk in Australian terrestrial vertebrates to the end of this century. These changes result in threatened terrestrial vertebrates becoming more concentrated within the current network of protected areas.

Our models indicate a sharp rise in extinctions after 2080 under the worst-case climate scenario (SSP5.85). This increase is driven by substantial warming later this century (from 2.4⍰°C in 2041– 2060 to 4.4⍰°C in 2081–2100 under SSP5.85^21^) and a marked decline in rainfall across southern Australia^22^. Overall, projected extinctions by 2100 range from 19–35 species under SSP1.26 (depending on dispersal assumptions) to 95–109 species under SSP5.85 (Table S7). This may be an underestimate given the increasing frequency and severity of extreme, climate-linked stochastic events such as the Australian 2019/2020 Black Summer megafires, which could result in the extirpation of threatened species long before changes in mean climatic conditions make areas inhospitable^23^. Furthermore, many more extinctions are likely across the four groups we analysed because the 577 species that we were unable to model are a non-random sample of Australia’s vertebrates, with smaller geographic range sizes and higher current threat status, on average (Figure S10). Of course, our predictions are associated with considerable uncertainty (*e*.*g*., modelling assumptions, dispersal capabilities, adaptive capacity) in addition to that captured by the range of scenarios we modelled. Nonetheless, these results have two important implications. First, the predicted surge in extinctions late this century underscores the importance of a long-term outlook for conservation planning to minimize biodiversity loss. Second, these results suggest that dozens of Australian vertebrate species could be saved from extinction by global-scale action to avoid the fossil-fuelled development scenario implied by SSP5.85.

Our models assume that in general, species that suffer the greatest loss of suitable climate area will be most likely to experience a severe contraction in their distribution. The extent to which this is true for any given species will depend on species-specific factors such as habitat specificity, the availability of suitable habitat into which a species can migrate, or changing spatial overlap with interacting species such as competitors, predators or pathogens (*e*.*g*., ^24^). Our focus in this study is on aggregate taxonomic and spatial patterns of extinction risk, which are likely to be relatively robust to such variation in species-specific responses to climate change. Nonetheless, our models also reveal particular species of concern, which are consistently predicted to become extinct under different climate-change scenarios (Table S7). Some are narrow-ranged habitat specialists that are already threatened (*e*.*g*., alpine bog-skink [*Pseudemoia cryodroma*], Howard River toadlet [*Uperoleia daviesae*] or northern hairy-nosed wombat [*Lasiorhinus krefftii*]). Others are not currently recognized as threatened (*e*.*g*., western false pipistrelle [*Flasistrellus mackenziei*], Karri frog [*Geocrinia rosea*] or silky mouse [*Pseudomys apodemoides*]), and hence unlikely to be considered a high priority for conservation, yet our models suggest they could become extinct, suggesting these species could be prioritized for research or early intervention.

While the IUCN threat classification scheme allows for future threats to be included in assessments under criterion D2^25^, there is no current provision for how already existing threatening processes might shift in extent and magnitude, and how this may change assessment outcomes. Our integrated framework represents the first attempt to model projected climate-driven changes in species distributions and projected changes in species threat status simultaneously and can identify which changes in threatening processes drive future predicted threat levels on a species-by-species basis (Figure S3). While this is no replacement for empirical, expert-led assessments, which require periodic updating as new threats emerge and species’ circumstances change, our approach provides a robust first step for over-the-horizon conservation planning.

Early work focused on predicted changes in species distributions in response to climate change suggested that the effectiveness of PA networks may decrease under climate change^9^. However, our results suggest that the representation effectiveness of the PA network is predicted to increase for threatened species (either currently or predicted to become threatened), but not for unthreatened species. This can be explained by the alignment of several factors. First, the prevailing pressure of climate change in Australia is expected to drive shifts in species distributions towards the south and east of the continent, so that the severe or complete loss of species’ suitable climate envelopes is expected to be most prevalent close to the hard boundary of the south-eastern coast (Figures S4 & S6). Second, the east and south-eastern coastal regions are parts of Australia with high concentrations of narrow-range endemic species, which are more likely to become threatened than the more broadly distributed species of inland and northern Australia, even if they are currently not threatened. Third, south-eastern Australia has a high density of PAs compared to other areas of the continent, and many of these are in upland regions that have traditionally been less likely to be developed for agriculture. Thus, a tendency for range expansions across inland, western, and northern Australia, combined with a tendency for range contractions and emergence of new threatened species in the southeast, leads to the contraction and concentration of threatened species richness hotspots in the PA-rich southeast. Similar patterns underlying observed or predicted increases in PA representation of species have been described in other countries in which areas of cooler climate are more likely to be the recipients of leading-edge expansion of species ranges from warmer areas^11–13,17,26–28^.

Our integrated framework predicts widespread changes in extinction risk across Australian terrestrial vertebrates, including an increased representation of threatened species within the current PA network. Some authors have interpreted similar findings in a positive light, representing an increase in the effectiveness of reserves under climate-driven range shifts. However, in our results, the increased representation of threatened species in PAs in the southeast is mirrored by sharp increases in extinctions in regions with fewer PAs across southern and northern Australia (Figure S4) and range contractions and shifts into areas with higher densities of PAs along Australia’s southeast (Figures S5-S6). These results suggest that patterns of extinction risk will be highly dynamic, so that the current PA system must also consider future risk. Australian government policy is to expand the coverage of the country’s PA system from the current ∼24.5% to 30% of the land area by 2030. This expansion should consider dynamic threat patterns and staged planning of new PA establishment in areas that may not yet be priorities but are predicted to become priority areas. Thus, future-proof conservation policy can be achieved by carefully incorporating principles of over-the-horizon threat assessment^4^. More broadly, our results highlight the importance of conservation actions not connected to the establishment of reserves, including global action on climate change and proactive local interventions such as invasive predator control or conservation fencing to secure species^29^.

## METHODS

### Data Collection – Species

We downloaded global shapefiles of the distributions of non-marine mammals and amphibians from the IUCN Red List^30^, of birds from BirdLife International^31^, and non-marine reptiles from the Global Assessment of Reptile Distributions (GARD) initiative v1.7^32^. These shapefiles were then cropped and filtered in R v4.5.0^33^ to only include polygons that overlap Australia in their native ranges, and, for birds, in their resident, breeding or non-breeding ranges. We removed 82 species of seabirds that are not known to breed in mainland Australia or Tasmania. Thus, our dataset did not include invasive, vagrant, or migratory species, but only species that have established native populations in Australia (n = 2,156).

We retrieved occurrence data for species from the Atlas of Living Australia (ALA^34^) using the *galah* package v2.1.1^35^. We filtered occurrences based on the ALA data quality profile (removing duplicate records, records with spatial or taxonomic issues, fossil and absence records, and possible outliers), with year of record > 1990 to ensure we include only relatively recent records with high spatial and taxonomic certainty. As an additional quality control, we then pruned out all occurrences that fell outside a 100 km buffer applied to each species’ range polygon to remove occurrences that were far outside known species ranges and are therefore likely to be erroneous (n = 2,132 species with occurrence records following data cleaning). We repeated the same process of downloading occurrence data (but without filtering within buffered range polygons) for five invasive species that are known to have large negative impacts on Australia’s terrestrial vertebrate fauna^36^, to be used as predictor variables for threat assessment: cane toads (*Rhinella marina*), cats (*Felis catus*), red foxes (*Vulpes vulpes*), camels (*Camelus dromedarius*), and European rabbits (*Oryctolagus cuniculus*).

We acquired current IUCN Red List assessments for all species from the IUCN Red List using the *taxize* package v0.10.0^37,38^. We manually updated the status of seven turtle species (*Chelodina longicollis, C. steindachneri, Elseya dentata, Emydura macquarii, E. subglossa, E. victoriae*, and *Myuchelys latisternum*) that do not have listings on the IUCN website but were assessed by the Turtle Taxonomy Working Group^39^. For species listed under Australia’s Environment Protection and Biodiversity Conservation Act 1999 (EPBC Act), we opted to use their EPBC listings (https://www.environment.gov.au/cgi-bin/sprat/public/publicthreatenedlist.pl) in lieu of their global IUCN assessment, as these are based on the same set of criteria as the IUCN Red List, but more accurately reflect the status of Australian populations of wider-ranging species (*e*.*g*., the curlew sandpiper, *Calidris ferruginea*, is globally listed as Near Threatened [NT], but nationally as Critically Endangered [CR]).

Finally, we collected data on body size of Australian terrestrial vertebrate species from published sources^40–56^ for the automated assessment procedure (see below) because size has been identified as a key predictor of extinction risk in vertebrates^57^. We used the natural logarithm of mean adult body mass (g) for all species to maintain comparability between our focal taxa.

### Data Collection – Environment

For the baseline current climatic conditions, which we used to model species ranges (see section: Species Distribution Modelling – training), we used historical climate data for 1970–2000 from WorldClim v2.1^58^. To encapsulate the mean, variation, and extremes in climatic variation, we chose the following WorldClim variables: BIO1 (annual mean temperature; °C), BIO4 (temperature seasonality; SD * 100), BIO10 (mean temperature of warmest quarter; °C), BIO11 (mean temperature of coldest quarter; °C), BIO12 (annual precipitation; mm), BIO15 (precipitation seasonality; coefficient of variation), BIO16 (precipitation of wettest quarter; mm), and BIO17 (precipitation of driest quarter; mm). These data were downloaded at a 2.5-minute resolution.

Animal species distributions are not determined solely by climate, but may also be limited by physical landscape features that influence distributions of plant species and vegetation types. We therefore included topographic and geological data in our SDMs (see section: Species Distribution Modelling – training). We obtained a Digital Elevation Model (DEM) at 2.5-minute resolution from WorldClim v2.1^58^, which was derived from the Shuttle Radar Topography Mission (SRTM) global elevation data. We obtained the following soil attributes from the Soil and Landscape Grid of Australia^59,60^: AWC (Available Water Capacity; %), BDW (Bulk Density – Whole earth; g/cm^3^), CLY (Clay content; %), pHc (pH - CaCl2), SLT (Silt content; %), SND (Sand content; %), SOC (organic carbon; %). We downloaded these data at a 3 second resolution at a depth of 0-5cm and then aggregated them by averaging across 2.5-minute resolution cells to match the resolution of the climatic data.

For anthropogenic conditions, which we used to model species threat status (see section: Species Automated Assessment), we downloaded baseline projections for 2010 of human population density (HPD) at 30 second resolution^61,62^, aggregated by averaging across 2.5-minute resolution cells, and land use (LU) at 30-minute resolution^63^. We filtered the LU raster to only include cells coded as “cropland_bioenergy”, “cropland_other”, and “built_up” and considered these as human-modified lands (note that this does not include the impacts of extensive rangeland grazing which covers a large amount of land area in Australia).

For future projections of climate (see section: Species Distribution Modelling – projections) and anthropogenic conditions (see section: Automated Assessment), we considered two Shared Socioeconomic Pathways (SSP1.26 and SSP5.85) under the HadGEM3-GC31-LL global circulation model, which has the best fit for Australia’s climatic patterns^64^, to capture uncertainty in projections of future climates, HPD, and LU. The SSPs represent upper and lower extremes in modelled trajectories in climate, demographics, economy, and land use, based on different climate change mitigation strategies. These range from Sustainability (low challenges to mitigation and adaptation; SSP1) to Fossil-Fuelled Development (high challenges to mitigation and low challenges to adaptation; SSP5). For each SSP we obtained projections at four 20-year intervals (2021–2040; 2041–2060; 2061–2080; 2081–2100) for HPD^61,62^, LU^63^, and downscaled Coupled Model Intercomparison Project Phase 6 (CMIP6)^65^ climate data from WorldClim v2.1^58^.

### Species Distribution Modelling – training

We used the filtered species occurrence data to generate SDMs for each species in the dataset with at least 10 occurrence records^66^ (n = 1,928; Table S7). To reduce spatial non-independence of presence points we performed spatial thinning by sampling a single point per grid cell in the environmental raster resolution (if more than one was present) using the `gridSample` function in the *dismo* package v1.3.5^67^.

Uneven sampling effort in presence points could lead to bias in modelled distributions if not accounted for when selecting background points^68,69^. To get an unbiased measure across space of how many samples are reported for each site, we selected background points while accounting for sampling bias by treating all occurrence data in that taxon (amphibians, birds, mammals, or reptiles) as the baseline background for the taxon. We performed spatial thinning^70^ on the background points by randomly selecting 10% of the records for mammals, reptiles, and amphibians, and 1% of the records for birds (due to the much higher number of bird occurrences in ALA).

For each species (n = 1,928), we fitted ensemble SDMs^71^ using the *biomod2* package v4.2-6-2^72^. We fitted five different algorithms using the `BIOMOD_modeling` function (Boosted Regression Trees [GBM], Generalised Additive Model [GAM], Generalised Linear Model [GLM], Maximum Entropy [MAXENT], and Random Forest [RF]) with the `bigboss` predefined model parameters. We assessed goodness-of-fit of individual models via spatially explicit block validation^73^ using the *blockCV* package v3.1.5^74^. For each species, we randomly divided the sampling space into hexagonal blocks, which were distributed among *k* folds. Block size (in m) was determined as the lower value of either the estimates spatial autocorrelation range (estimated using the `cv_spatial_autocor` function) or the square root of the area of the species’ range shapefile. The number of folds *k* was then determined in a stepwise fashion, starting with 4 and attempting to generate spatial blocks using the `cv_spatial` function, and decreasing *k* by 1 until spatial blocks were successfully generated. If no blocks were successfully generated with *k*=2, SDM fitting for this species was skipped and it was excluded in downstream analyses (n = 4 species, all reptiles). After individual models were fit, an ensemble model was generated by calculating a weighted mean of all models, weighing models by their calculated True Skill Statistic (TSS; a measure of model accuracy calculated from the sum of sensitivity [true positive rate] and specificity [true negative rate]), after filtering out individual models with negative values of TSS, which are indicative of predictions no better than random chance. Five species (four reptiles, one bird) had no individual models with positive TSS values. Ensemble model fitting failed for an additional four species (one amphibian, one mammal, two reptiles) for unknown reasons (error in running the `BIOMOD_modeling` function). All nine were excluded from downstream analyses. Ensemble models were constructed for a total of 1,914 species (Table S8).

### Species Distribution Modelling – projections

For each modelled species, we used the ensemble model to make predictions of climatic suitability across Australia for current conditions and for future conditions in each timestep (up to 2040, 2060, 2080, 2100) under the two SSPs examined. We converted predicted suitability into binary presence-absence predictions using a threshold value, which was selected for each species by maximising its TSS using the ‘bm_FindOptimStat’ function. Summaries of SDMs for each species, including sample sizes, evaluation metrics and threshold values, can be found in Table S9.

We used the binary predicted occurrence layers from the SDMs as proxies for predicted extent of occurrence in future timesteps. However, treating the entire range above-the-threshold as future extent of occurrence does not take into account species’ differing abilities to disperse to track shifting climatic conditions. In reality, climatically “suitable” areas that are too far to reach will be outside a future extent of occurrence. Therefore, for future predictions of presence, we established an additional pipeline to make sure that range shifts were possible within reasonable dispersal limits, defined by sequentially applying dispersal buffers to suitability layers in each timestep. In order to be counted as presence cells in a future extent of occurrence map, cells that exceeded the presence threshold also needed to be within the limits of dispersal buffer applied to the previous timestep.

Due to limited data for dispersal capabilities for most of Australia terrestrial vertebrate species, we bracketed uncertainty by considering three different dispersal scenarios to generate buffers. In the first (“none”), the dispersal buffer was set to 0, disallowing any range expansions in subsequent timesteps, reflective of species that cannot track range shifts due to limited intrinsic dispersal capabilities or hard geographic barriers. In the second (“limited”), we estimated a fixed dispersal limit for all species based on available mammal data. We calculated the dispersal buffer by estimating maximal dispersal rate per generation for Australian mammals from body mass, trophic level, and home range size^75^ using published power laws^76^. We then divided the maximal rate by generation length (using year at first birth as a surrogate^75^) to arrive at a per year dispersal rate of, on average, ∼7km^77^, or ∼140km over a 20-year interval. We then rounded this value to 150km and applied it as the dispersal buffer. Since this value is averaged over several mammal species (the only taxon for which the required data were available), this likely represents an underestimate of dispersal capability for highly mobile animals (*e*.*g*., large mammals, bats, birds), and an overestimate for species with low mobility (*e*.*g*., legless lizards, frogs). In the third dispersal scenario (“unlimited”), the dispersal buffers were designated as the entire landmass in which species are present, only disallowing overwater dispersal.

The same SDM procedures were repeated for the five invasive species, treating the entire pool of occurrence data for the invasive species as the background points, to estimate current and future range sizes for each of the invasive species. We then calculated for each native species, across its current and predicted future extents of occurrence, the range size (km^2^), percentage of range overlap with cells that have over 50% human-modified land, percentage of range overlap with cells that have over 100/km^2^ human population density, mean % of human-modified land averaged across the entire range, mean human population density averaged across the entire range (cut-off values for the human population density and land use features as in ^5^), and percentage of range overlap with each of the five invasive species. These values were then used as features for the automated assessment algorithm.

### Automated Assessment of Threat Status

We used a machine learning-based automated assessment method originally developed to infer current threat status for Data Deficient (DD) or Not Evaluated (NE) reptile species^6^, and extended it to allow it to predict IUCN threat categories (LC: Least Concern; NT: Near Threatened; VU: Vulnerable; EN: Endangered; CR: Critically Endangered) under future climate change scenarios. Due to a low sample size of CR species in our dataset (Table S8), we combined EN and CR into a single category. Our automated assessment uses eXtreme Gradient Boosting (XGBoost^78^), an efficient and highly accurate supervised machine learning algorithm^79^. The approach relies on training a model to perform hierarchical binary classification tasks in a decision tree: first separating threatened (VU, EN/CR) from non-threatened (LC, NT) species; then, separating LC from NT species; then, separating EN/CR from VU species (see detailed flowchart in Figure 5 in ^6^). Each binary classification task involves hyperparameter tuning and model fitting, using nested cross-validation to test prediction accuracy (see below). In addition to species omitted due to failure to fit SDMs, we did not train the model on species whose predicted ranges greatly exceeded range shapefiles (≥ 3 times larger or smaller; n = 226). This is to reduce errors in classification due to inflated range sizes, because range size is one of the key predictors for IUCN threat level (based on criterion B). Although these species were excluded from model training, we used the XGboost model to predict their threat levels, and these were used in the downstream analyses. In total, we trained XGBoost models on 1631 species, 76% of Australia’s terrestrial vertebrate fauna (Table S8). We note that species omitted from training for automated assessment (n = 525) are disproportionally threatened and narrow-ranged (Figure S10), so our assessments likely represent a conservative estimate of future threat levels.

To predict threat status for each species, we included seven features for each species: mass (g), range size (km^2^), percentage of range overlap with cells that have over 50% human-modified land, percentage of range overlap with cells that have over 100/km^2^ human population density, mean percentage of human-modified land averaged across the entire range, mean human population density averaged across the entire range, and percentage of range overlap with each of the five invasive species. Given the relatively small number of features, we did not perform feature selection as in ^6^, so all features are included in the model to predict species threat.

In addition to these features, we also considered which higher taxon (class) each species belongs to, and which biome each species’ distribution is predominantly found in. To assign species to biomes we overlapped species ranges with shapefiles of terrestrial ecoregions in Australia^80^, and assigned each species to the biome in which the highest proportion of its distribution lies. We then partitioned all species into three biome categories, to ensure enough species in each combination of biome, class, and threat level. The three biome categories are: tropical (Tropical and Subtropical Moist Broadleaf Forests; Tropical and Subtropical Grasslands, Savannas, and Shrublands), temperate (Temperate Broadleaf and Mixed Forests; Temperate grasslands, Savannas, and Shrublands; Montane Grasslands and Shrublands), and dry (Mediterranean Forests, Woodlands, and Scrub; Deserts and Xeric Shrublands). Interactions among features, classes, and biomes were fully allowed in model training, since threatening processes in Australia vary regionally across habitats^81^ and classes^82,83^.

We trained the XGBoost models using the *xgboost* package v1.7.10.1^84^. We tuned the following hyperparameters: learning rate, maximum tree depth, minimum child weight, row sampling, column sampling, weight balancing, and the regularisation parameters γ, α, and λ, by randomly sampling 10,000 different values for each parameter. Given the size of our dataset we applied nested cross-validation (rather than setting aside a proportion of the dataset as validation data and test data as in ^6^): the dataset was first split into *N* folds; each fold was used as test data with the other *N*-1 folds used as training data to fit models. These *N*-1 folds were further split into *n* folds; each of these *n* folds was used as validation data with the other *n*-1 folds used as training data to tune hyperparameters. This nested cross-validation ensured that training, validation, and test data have no overlap.

We created custom code to ensure that different folds have roughly similar numbers of species in each combination of biome, class, and threat level. When the number of species in a combination was smaller than *N*, we set aside these species to the test data, so that these underrepresented species will not cause model instability, and the prediction accuracy can reflect how much information the other species carry on the threatening process of these underrepresented species. For the dataset to separate threatened from non-threatened species, we set *N*=*n*=5. The datasets to separate LC from NT species and to separate CR/EN and VU species were much smaller, so for them we set *N*=5 and *n*=3 (LC from NT) and *N*=4 and *n*=2 (CR/EN from VU).

At the end of the nested cross-validation, the means of the best fit hyperparameters for each of the *N* folds were used as the hyperparameters for the final model fitting using all the data. The average of the prediction accuracy of each of the *N* test data was used as the prediction accuracy of the final model (Table S1). To assess model stability, we repeated the whole procedure three times, and all replicates gave similar fitted models and prediction accuracy.

We then assessed relative feature importance for prediction in the final model, ranking features by estimated gain (fractional contribution of each feature to the model based on the total gain of this feature’s splits). This represents the contribution of each feature to model predictions, representing its relative importance in assigning species to threat categories. Finally, we used the final model to predict IUCN threat levels. To set a baseline against which to compare future predicted threat levels, we use the same method to predict current threat levels for all 1,914 species with modelled distributions (1,631 species used to train the model and 283 species that were either not assessed [NE], assessed as data deficient [DD], or were dropped due to having predicted ranges much larger than shapefiles; see above). This is because current IUCN threat levels are determined empirically by expert assessors based on additional information (such as observed rates of population change) so would not be directly comparable to model-based inferences of threat level. We made predictions for future threat status under all examined climate change and dispersal scenarios using the calculated values of the features for estimated future ranges for all 1,914 species. We classified a species as extinct if it was not predicted to occur in any cell in the corresponding timestep, after accounting for dispersal as described above.

### Analysis of changing taxonomic and spatial patterns of threat

To explore whether species are predicted to move through the IUCN Red List sequentially (*i*.*e*., increase in threat through time), we fitted an ordered logistic regression to the predicted IUCN threat status using the *MASS* package v7.3.65^85^. The response variable was the predicted IUCN threat status in 2040, 2060, 2080 and 2100 as inferred from automated assessment. The independent variables were the IUCN threat status in the previous timestep, the modelling scenario (combination of SSP and dispersal scenario; *e*.*g*., SSP2.1 no dispersal), and the taxonomic Class.

To explore general patterns across species in the way ranges are predicted to shift with climate change, we calculated the following three metrics for predicted ranges in each time step and under each climate change and dispersal scenario:

1. Direction of range shift: We calculated the range centroid for the predicted range of each species at each timestep. We then calculated the azimuth of each centroid compared to the centroid in the previous time step for all predicted ranges in 2040, 2060, 2080, and 2100, with 0° being due north.
2. Distance shifted: We calculated the Euclidean distance in km between each centroid to the centroid in the previous time step for all predicted ranges in 2040, 2060, 2080, and 2100, after projecting all ranges to an equal area Australia Albers projection (EPSG:3577).
3. Magnitude of range contraction/expansion: We calculated the area in km^2^ of all predicted ranges after projecting to an equal area Australia Albers projection (EPSG:3577). We then calculated log(area in each time step /area in the previous timestep) to generate a symmetrical index where negative values indicate range contraction and positive values indicate range expansion.

We examined how spatial patterns of threatened species richness change across Australia with time. For each time step and climate change scenario we overlayed all predicted ranges of species predicted to be threatened in that time step on a map of Australia in an equal-area Australian Albers projection (EPSG:3577) and tallied the number of threatened species per 50x50km cell. Conservation planning is often based on identifying areas with concentrations of threatened species from different higher-level taxa (*e*.*g*., ^7^), so the congruence of patterns of threatened species richness— the degree to which an area that contains many threatened species for one taxon is also expected to contain many threatened species of other taxa—may have important implications for the efficiency of planning for protected areas to maximize biodiversity protection. We assessed congruence in threatened species richness by calculating spatially corrected Pearson’s correlation coefficients (Tjostheim’s coefficient^86,87^) using the `cor.spatial` function from the *SpatialPack* package v0.4.1^88^. We calculated congruence between threatened species richness of each pair of terrestrial vertebrate classes in each time step and climate change scenario after omitting double-zero cells. We used an ANCOVA to assess whether congruence (Tjostheim’s coefficient) between different taxa changes over years.

Finally, we defined hotspots of threatened species richness for each vertebrate class as the cells in the top 10^th^ percentile of threatened species richness. We downloaded a layer of protected areas in Australia from Collaborative Australian Protected Area Database (CAPAD^89^). We calculated overlap of threatened species hotspots with current protected areas as the proportion of hotspots cells whose centroids fall within protected areas. We then compared the overlap of threatened species hotspots with protected areas in different timesteps and climate change scenarios to assess the adequacy of Australia’s current network of protected areas for protecting species in the future as their ranges and threat levels change due to climate change.

## ACKNOWLEDGEMENTS

This research was funded by the College of Science, Australian National University. BCS was supported by The Australian Research Council (ARC) through a Discovery Early Career Research Award (DE200100121) and a Future Fellowship (FT250100113).

## AUTHOR CONTRIBUTIONS

Conceptualization – AS, MC, LB, XH, BCS; Data curation – AS; Software – AS, XH; Formal analysis – AS; Funding acquisition – MC, LB, XH, BCS; Investigation – AS; Visualization – AS; Writing – original draft – AS, MC, LB, XH, BSC; Writing – review & editing – AS, MC, LB, XH, BSC.

## References

1. Johnson, C. N. et al. Biodiversity losses and conservation responses in the Anthropocene. Science (1979). 356, 270–275 (2017).

2. Díaz, S. et al. Pervasive human-driven decline of life on Earth points to the need for transformative change. Science (1979). 366, eaax3100 (2019).

3. Foden, W. B. et al. Climate change vulnerability assessment of species. WIREs Climate Change 10, (2019).

4. Cardillo, M., Scheele, B. C. & Tulloch, A. I. T. Forecasting extinction risk for future-proof conservation decisions. Trends Ecol. Evol. 41, 112–119 (2026).

5. Cardillo, M., Skeels, A. & Dinnage, R. Priorities for conserving the world’s terrestrial mammals based on over-the-horizon extinction risk. Current Biology 33, 1381–1388 (2023).

6. Caetano, G. H. de O. et al. Automated assessment reveals that the extinction risk of reptiles is widely underestimated across space and phylogeny. PLoS Biol. 20, e3001544 (2022).

7. Grenyer, R. et al. Global distribution and conservation of rare and threatened vertebrates. Nature 444, 93–96 (2006).

8. Myers, N., Mittermeier, R. A., Mittermeier, C. G., Da Fonseca, G. A. B. & Kent, J. Biodiversity hotspots for conservation priorities. Nature 403, 853–858 (2000).

9. Araújo, M. B., Cabeza, M., Thuiller, W., Hannah, L. & Williams, P. H. Would climate change drive species out of reserves? An assessment of existing reserve-selection methods. Glob. Chang. Biol. 10, 1618–1626 (2004).

10. Hannah, L. et al. Protected area needs in a changing climate. Front. Ecol. Environ. 5, 131–138 (2007).

11. Thomas, C. D. & Gillingham, P. K. The performance of protected areas for biodiversity under climate change. Biological Journal of the Linnean Society 115, 718–730 (2015).

12. Thomas, C. D. et al. Protected areas facilitate species’ range expansions. Proceedings of the National Academy of Sciences 109, 14063–14068 (2012).

13. Lehikoinen, P., Santangeli, A., Jaatinen, K., Rajasärkkä, A. & Lehikoinen, A. Protected areas act as a buffer against detrimental effects of climate change—Evidence from large-scale,long-term abundance data. Glob. Chang. Biol. 25, 304–313 (2019).

14. Coldrey, K. M. & Turpie, J. K. The future representativeness of Madagascar’s protected area network in the face of climate change. Afr. J. Ecol. 59, 253–263 (2021).

15. Popescu, V. D., Rozylowicz, L., Cogălniceanu, D., Niculae, I. M. & Cucu, A. L. Moving into protected areas? Setting conservation priorities for Romanian reptiles and amphibians at risk from climate change. PLoS One 8, e79330 (2013).

16. Critchlow, R. et al. Multi-taxa spatial conservation planning reveals similar priorities between taxa and improved protected area representation with climate change. Biodivers. Conserv. 31, 683–702 (2022).

17. Hole, D. G. et al. Projected impacts of climate change on a continent-wide protected area network. Ecol. Lett. 12, 420–431 (2009).

18. Mi, C. et al. Global Protected Areas as refuges for amphibians and reptiles under climate change. Nat. Commun. 14, 1389 (2023).

19. Peng, S. et al. Incorporating global change reveals extinction risk beyond the current Red List. Current Biology 33, 3669–3678 (2023).

20. Visconti, P. et al. Future hotspots of terrestrial mammal loss. Philosophical Transactions of the Royal Society B: Biological Sciences 366, 2702–2963 (2011).

21. Calvin, K. et al. IPCC, 2023: Climate Change 2023: Synthesis Report. Contribution of Working Groups I, II and III to the Sixth Assessment Report of the Intergovernmental Panel on Climate Change [Core Writing Team, H. Lee and J. Romero (Eds.)]. IPCC, Geneva, Switzerland. (2023) doi:10.59327/IPCC/AR6-9789291691647.

22. CSIRO & Bureau of Meteorology. Climate Change in Australia. Preprint at http://www.climatechangeinaustralia.gov.au/ (2023).

23. Heard, G. W. et al. Drought, fire, and rainforest endemics: A case study of two threatened frogs impacted by Australia’s “Black Summer”. Ecol. Evol. 13, e10069 (2023).

24. Sopniewski, J., Scheele, B. C. & Cardillo, M. Predicting the distribution of Australian frogs and their overlap with Batrachochytrium dendrobatidis under climate change. Divers. Distrib. 28, 1255–1268 (2022).

25. IUCN. IUCN Red List Categories and Criteria: Version 3.1. (International Union for Conservation of Nature, Gland, Switzerland, 2012).

26. Anderson, B. J. & Ohlemüller, R. Climate change and protected areas: how well do British rare bryophytes fare? in Bryophyte Ecology and Climate Change (eds. Tuba, Z., Slack, N. G. & Stark, L. R.) 409–425 (Cambridge University Press, Cambridge, UK, 2010).

27. Thuiller, W. et al. Are different facets of plant diversity well protected against climate and land cover changes? A test study in the French Alps. Ecography 37, 1254–1266 (2014).

28. Virkkala, R., Pöyry, J., Heikkinen, R. K., Lehikoinen, A. & Valkama, J. Protected areas alleviate climate change effects on northern bird species of conservation concern. Ecol. Evol. 4, 2991–3003 (2014).

29. Scheele, B. C. et al. How to improve threatened species management: An Australian perspective. J. Environ. Manage. 223, 668–675 (2018).

30. IUCN. The IUCN Red List of Threatened Species. Version 6.2. https://www.iucnredlist.org. Downloaded on 02 July 2022. Preprint at (2019).

31. BirdLife International & Handbook of the Birds of the World. Bird species distribution maps of the world. Version 2022.2. Preprint at (2022).

32. Roll, U. et al. The global distribution of tetrapods reveals a need for targeted reptile conservation. Nat. Ecol. Evol. 1, 1677–1682 (2017).

33. R Core Team. R: a language and environment for statistical computing. Preprint at (2025).

34. Belbin, L., Wallis, E., Hobern, D. & Zerger, A. The Atlas of Living Australia: History, current state and future directions. Biodivers. Data J. 9, e65023 (2021).

35. Westgate, M., Kellie, D., Stevenson, M. & Newman, P. galah: Biodiversity Data from the GBIF Node Network. R package version 2.1.1. Preprint at https://CRAN.R-project.org/package=galah (2025).

36. Johnson, C. N. Australia’s Mammal Extinctions: A 50,000-Year History. (Cambridge University Press, Cambridge, UK, 2006).

37. Chamberlain, S. & Szöcs, E. taxize - taxonomic search and retrieval in R. F1000Res. 2, 191 (2013).

38. Chamberlain, S. et al. taxize: Taxonomic information from around the web. Preprint at https://github.com/ropensci/taxize (2020).

39. Turtle Taxonomy Working Group. Turtles of the World: Annotated Checklist and Atlas of Taxonomy, Synonymy, Distribution, and Conservation Status (9th Ed.). in Conservation Biology of Freshwater Turtles and Tortoises: A Compilation Project of the IUCN/SSC Tortoise and Freshwater Turtle Specialist Group (eds. Rhodin, A. G. J. et al.) 1–472 (Chelonian Research Monographs 8, 2021).

40. Cooney, C. R. & Thomas, G. H. Heterogeneous relationships between rates of speciation and body size evolution across vertebrate clades. Nat. Ecol. Evol. 5, 101–110 (2021).

41. Itescu, Y. A biogeographic perspective on turtle evolution. (2012).

42. Thomson, S. & Georges, A. A new species of freshwater turtle of the genus Elseya (Testudinata: Pleurodira: Chelidae) from the Northern Territory of Australia. Zootaxa 4061, 18–28 (2016).

43. Oliveira, B. F., São-Pedro, V. A., Santos-Barrera, G., Penone, C. & Costa, G. C. AmphiBIO, a global database for amphibian ecological traits. Sci. Data 4, 1–7 (2017).

44. Slavenko, A. et al. Evolution of sexual size dimorphism in tetrapods is driven by varying patterns of sex-specific selection on size. Nat. Ecol. Evol. 9, 464–473 (2025).

45. Meiri, S. Traits of the lizards of the world: variation around a successful evolutionary design. Global Ecology and Biogeography 27, 1168–1172 (2018).

46. Storr, G. M. Geographic races of the agamid lizard Amphibolurus caudicinctus. J. R. Soc. West. Aust. 50, 49–56 (1967).

47. Fuery, C. J., Withers, P. C., Hobbs, A. A. & Guppy, M. The role of protein synthesis during metabolic depression in the Australian desert frog Neobatrachus centralis. Comp. Biochem. Physiol. A Mol. Integr. Physiol. 119, 469–476 (1998).

48. Iverson, J. B., Ennen, J. & Lovich, J. Life-history and ecology data for the turtles of the world. Chelonian Conservation and Biology 24, (2025).

49. Meiri, S. <scp>SquamBase</scp> —A database of squamate (Reptilia: Squamata) traits. Global Ecology and Biogeography 33, e13812 (2024).

50. Parish, S., Richards, G. & Hall, L. A Natural History of Australian Bats⍰: Working the Night Shift. (CSIRO Publishing, Melbourne, VIC, Australia, 2012).

51. Parnaby, H. E. A taxonomic review of Australian Greater Long-eared Bats previously known as Nyctophilus timoriensis (Chiroptera: Vespertilionidae) and some associated taxa. Australian Zoologist 35, 39–81 (2009).

52. Reardon, T. B. et al. A molecular and morphological investigation of species boundaries and phylogenetic relationships in Australian free-tailed bats Mormopterus (Chiroptera: Molossidae). Aust. J. Zool. 62, 109–136 (2014).

53. Reardon, T. B., Adams, M., McKenzie, N. L. & Jenkins, P. A new species of Australian freetail bat Mormopterus eleryi sp. nov. (Chiroptera: Molossidae) and a taxonomic reappraisal of M. norfolkensis (Gray). Zootaxa 1875, 1–31 (2008).

54. Richards, S. J. & Alford, R. A. Structure and dynamics of a rainforest frog (Litoria genimaculata) population in northern Queensland. Aust. J. Zool. 53, 229–236 (2005).

55. Silla, A. J., McFadden, M. & Byrne, P. G. Hormone-induced spawning of the critically endangered northern corroboree frog Pseudophryne pengilleyi. Reproduction, Ferility and Development 30, 1352–1358 (2018).

56. Slavenko, A., Tallowin, O. J. S., Itescu, Y., Raia, P. & Meiri, S. Late Quaternary reptile extinctions: size matters, insularity dominates. Global Ecology and Biogeography 25, (2016).

57. Cardillo, M. & Bromham, L. Body size and risk of extinction in Australian mammals. Conservation Biology 15, 1435–1440 (2001).

58. Fick, S. E. & Hijmans, R. J. WorldClim 2: new 1km spatial resolution climate surfaces for global land areas. International Journal of Climatology 37, 4302–4315 (2017).

59. Viscarra Rossel, R. A. et al. Soil and Landscape Grid National Soil Attribute Maps (3” resolution) - Release 1. v6. CSIRO. Data Collection. https://doi.org/10.4225/08/546ED604ADD8A (2014) doi:10.4225/08/546ED604ADD8A.

60. Malone, B. & Searle, R. Soil and Landscape Grid National Soil Attribute Maps (3” resolution) - Release 2. v4. CSIRO. Data Collection. (2022).

61. Jones, B. & O’Neill, B. C. Spatially explicit global population scenarios consistent with the Shared Socioeconomic Pathways. Environmental Research Letters 11, 84003 (2016).

62. Gao, J. Global 1-Km Downscaled Population Base Year and Projection Grids Based on the Shared Socioeconomic Pathways, Revision 01. (NASA Socioeconomic Data and Applications Center (SEDAC), Palisades, NY, 2020).

63. Fujimori, S., Hasegawa, T., Ito, A., Takahashi, K. & Masui, T. Gridded emissions and land-use data for 2005-2100 under diverse socioeconomic and climate mitigation scenarios. Sci. Data 5, 180210 (2018).

64. Adhikari, R. K., Yilmaz, A. G., Mainali, B. & Dyson, P. Performance evaluation of CMIP6 models for application to hydrological modelling studies – A case study of Australia. Science of The Total Environment 945, 174015 (2024).

65. Eyring, V. et al. Overview of the Coupled Model Intercomparison Project Phase 6 (CMIP6) experimental design and organization. Geosci. Model Dev. 9, 1937–1958 (2016).

66. Santini, L., Benítez-López, A., Maiorano, L., Čengić, M. & Huijbregts, M. A. J. Assessing the reliability of species distribution projections in climate change research. Divers. Distrib. 27, 1035–1050 (2021).

67. Hijmans, R. J., Phillips, S. & Elith, J. dismo: Species Distribution Modeling. Preprint at (2021).

68. Phillips, S. J. et al. Sample selection bias and presence-only distribution models: implications for background and pseudo-absence data. Ecological Applications 19, 181–197 (2009).

69. Elith, J. et al. A statistical explanation of MaxEnt for ecologists. Divers. Distrib. 17, 43–57 (2011).

70. Kramer-Schadt, S. et al. The importance of correcting for sampling bias in MaxEnt species distribution models. Divers. Distrib. 19, 1366–1379 (2013).

71. Araújo, M. B. & New, M. Ensemble forecasting of species distributions. Trends Ecol. Evol. 22, 42–47 (2007).

72. Thuiller, W. et al. biomod2: Ensemble Platform for Species Distribution Modeling. R package version 4. 2-6-2. Preprint at (2025).

73. Roberts, D. R. et al. Cross-validation strategies for data with temporal, spatial, hierarchical, or phylogenetic structure. Ecography 40, 913–929 (2017).

74. Valavi, R., Elith, J., Lahoz-Monfort, J. J. & Guillera-Arroita, G. <scp>block</scp><scp>CV</scp>1: An <scp>r</scp> package for generating spatially or environmentally separated folds for k-fold cross-validation of species distribution models. Methods Ecol. Evol. 10, 225–232 (2019).

75. Jones, K. E. et al. PanTHERIA: a species-level database of life history, ecology, and geography of extant and recently extinct mammals. Ecology 90, 2648 (2009).

76. Santini, L. et al. Ecological correlates of dispersal distance in terrestrial mammals. Hystrix: the Italian Journal of Mammalogy 24, 181–186 (2013).

77. Schloss, C. A., Nuñez, T. A. & Lawler, J. J. Dispersal will limit ability of mammals to track climate change in the Western Hemisphere. Proceedings of the National Academy of Sciences 109, 8606–8611 (2012).

78. Chen, T. & Guestrin, C. XGBoost: a scalable tree boosting system. in Proceedings of the 22nd ACM SIGKDD International Conference on Knowledge Discovery and Data Mining 785–794 (ACM, New York, NY, USA, 2016). doi:10.1145/2939672.2939785.

79. Nielsen, D. Tree boosting with XGBoost - why does XGBoost win ‘every’ machine learning competition? (Norwegian University of Science and Technology, Trondheim, Norway, 2016).

80. Olson, D. M. et al. Terrestrial ecoregions of the world: a new map of life on Earth. Bioscience 51, 933–938 (2001).

81. Cresswell, I. D., Janke, T. & Johnston, E. L. Australia State of the Environment 2021: Overview, Independent Report to the Australian Government Minister for the Environment, Commonwealth of Australia, Canberra. (2021).

82. Ducatez, S. & Shine, R. Drivers of extinction risk in terrestrial vertebrates. Conserv. Lett. 10, 186–194 (2017).

83. Harfoot, M. B. J. et al. Using the IUCN Red List to map threats to terrestrial vertebrates at global scale. Nat. Ecol. Evol. 5, 1510–1519 (2021).

84. Chen, T. et al. xgboost: Extreme Gradient Boosting. R package version 1.7.10.1. Preprint at 10.32614/CRAN.package.xgboost (2025).

85. Venables, W. N. & Ripley, B. D. Modern Applied Statistics with S. (Springer, New York, USA, 2002).

86. Tjøstheim, D. A measure of association for spatial variables. Biometrika 65, 109–114 (1978).

87. Hubert, L. & Golledge, R. G. Measuring association between spatially defined variables: Tjøstheim’s coefficient index and some extensions. Geogr. Anal. 14, 273–278 (1982).

88. Vallejos, R., Osorio, F. & Bevilacqua, M. Spatial Relationships Between Two Georeferenced Variables: With Applications in R. (Springer, New York, NY, USA, 2020).

89. Commonwealth of Australia. Collaborative Australian Protected Areas Database (CAPAD) 2020. (2021).

