## Supplementary material for "Over-the-horizon extinction risk assessment reveals rapidly shifting geographic and taxonomic priorities for conservation": Figures S1-S10; Tables S1-S6; Table S8

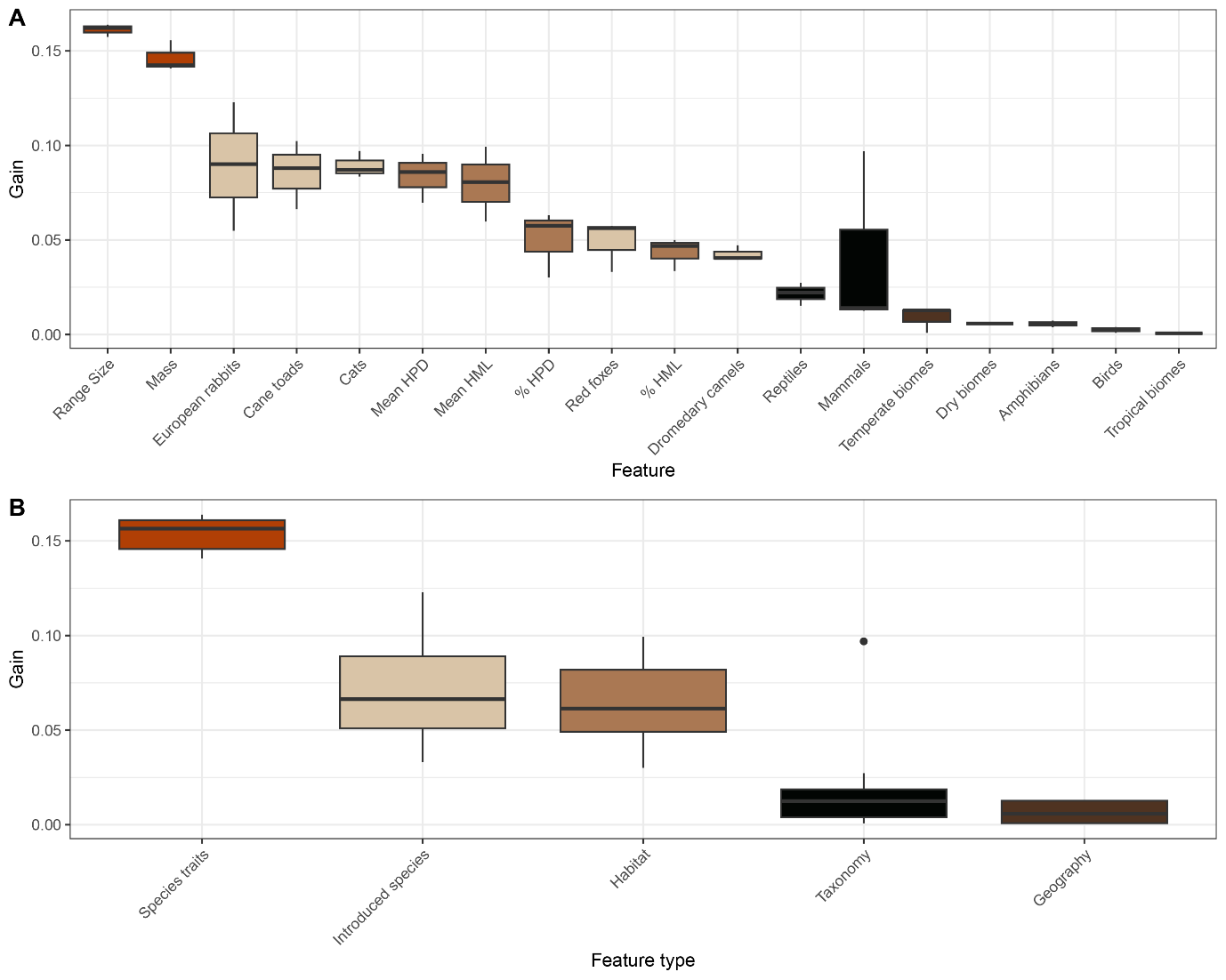

**Figure S1. Feature importance in the automated assessment algorithm**. Box plots showing the distribution of gain values for different features in all stages of the XGBoost automated assessment algorithm. HPD and HML refer to human population density and human-modified land, respectively. Mean of those values refers to the average across species’ ranges, and % refers to the proportion of species’ ranges overlapping areas with human population densities ≥ 100/km^2^ and areas that are ≥ 50% human modified, respectively. Features are ordered left to right by decreasing median gain (across all taxa). (A) Gain values for each individual feature across all XGBoost stages. (B) Gain values for categories of features, species traits (range size and mass), habitat (HPD and HML), introduced species, taxonomy (classes), and geography (biomes).

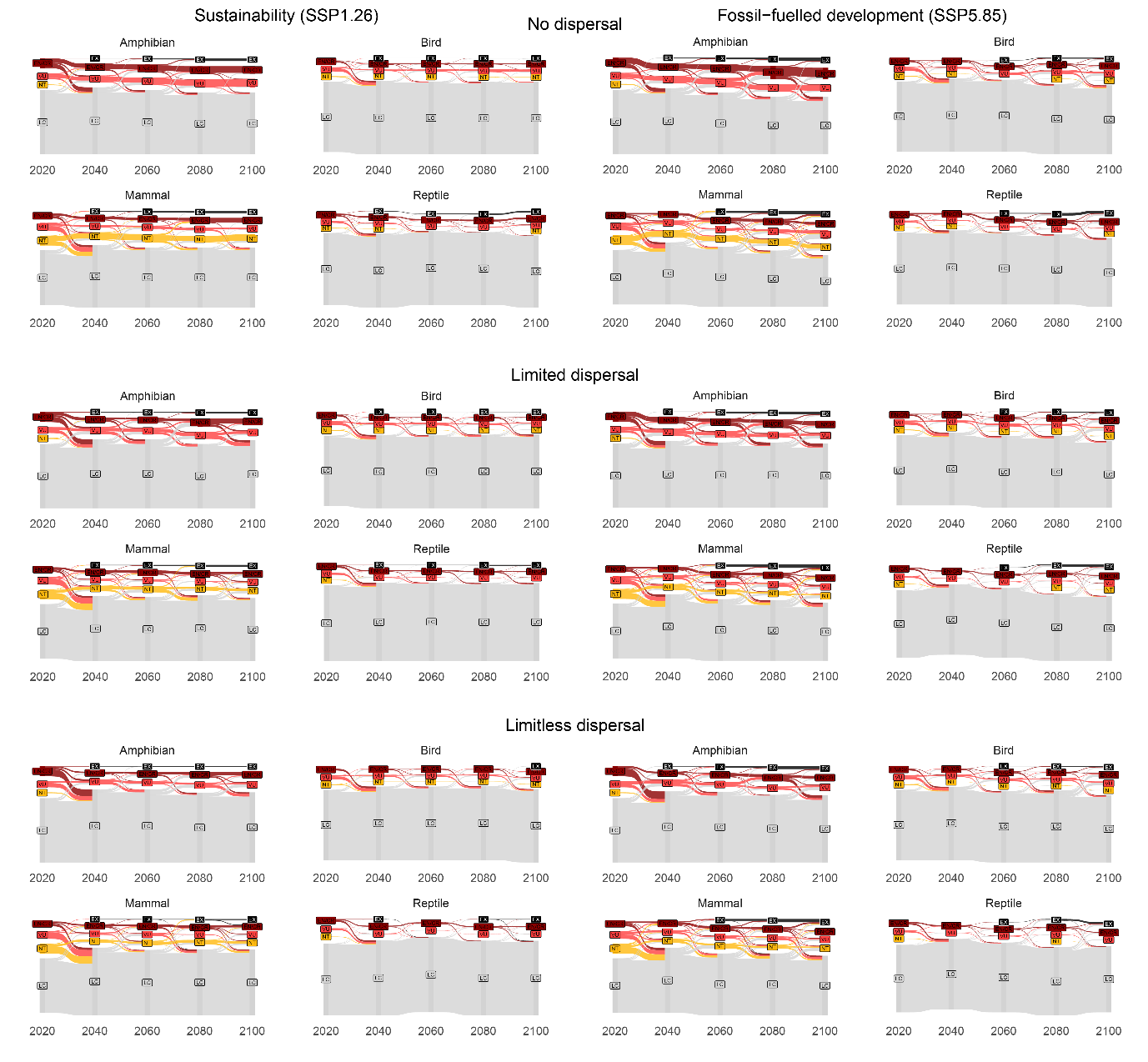

**Figure S2. Movement of species between IUCN categories.** Sankey plots showing shifts of species between different IUCN threat categories in 20-year timesteps under all examined climate change and dispersal scenarios.

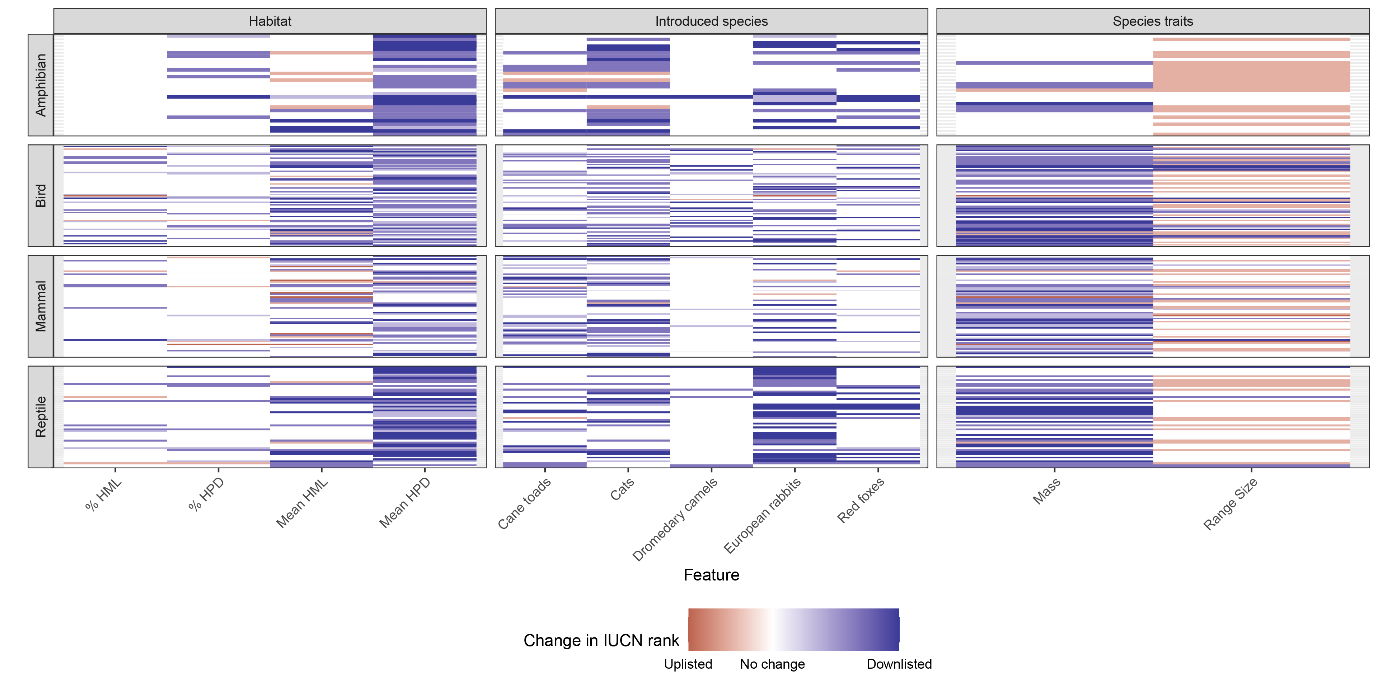

**Figure S3. Impact of features on individual future threat assessments.** Changes in predicted IUCN threat levels at 2100 based on a leave-one-out sensitivity analysis, where an individual feature is omitted from the prediction. The colours represent the change in the predicted assessment compared to the original prediction, which includes all features. Blue colours represent a lower relative threat level compared to the original prediction (*e.g.*, LC vs NT), whereas red colours represent a higher relative threat level (*e.g.*, EN/CR vs VU).

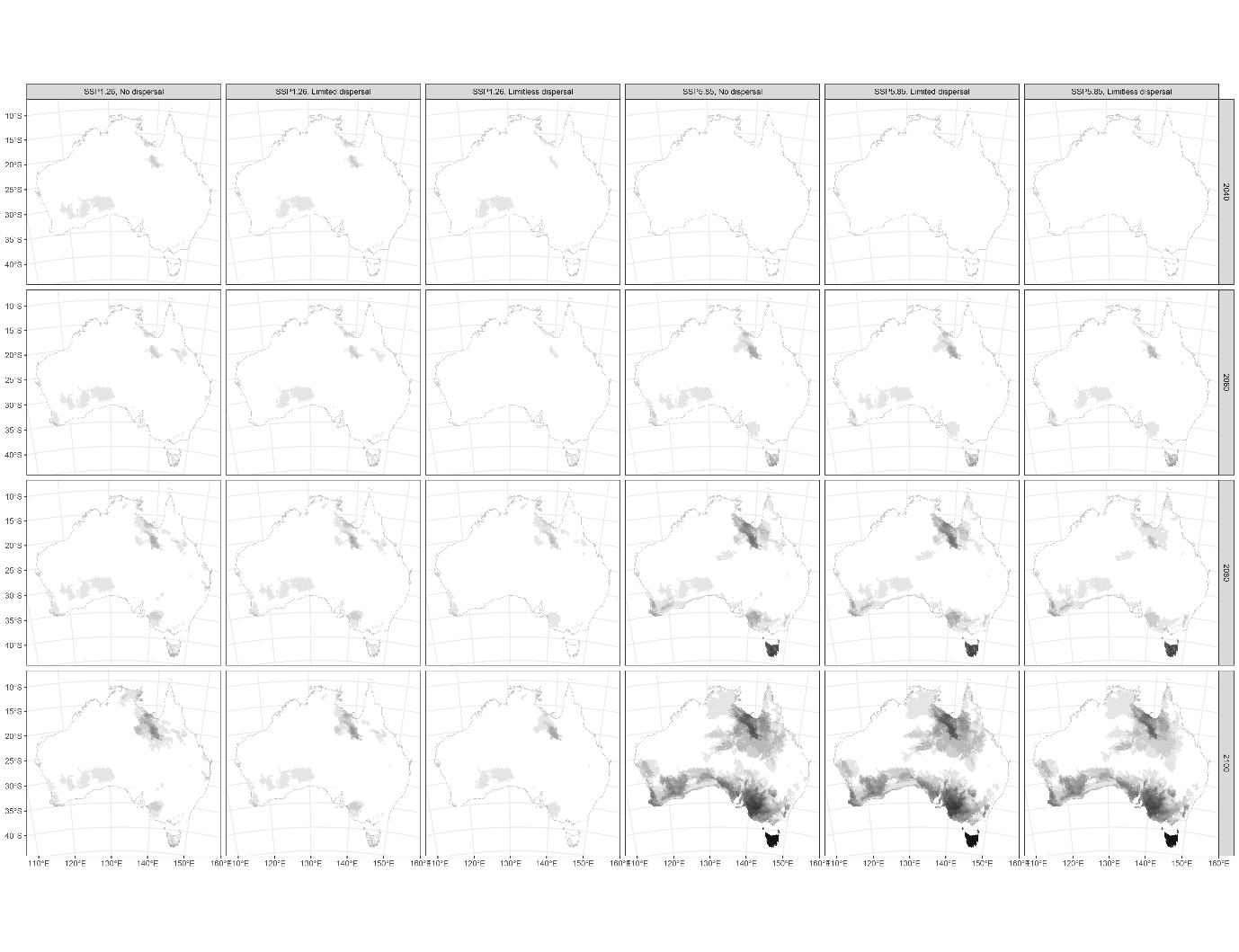

**Figure S4. Extinctions by the end of the century**. Maps showing the current distributions of species predicted to go extinct by the end of each timestep under all examined scenarios of climate change and dispersal. Darker shades represent more species predicted to go extinct.

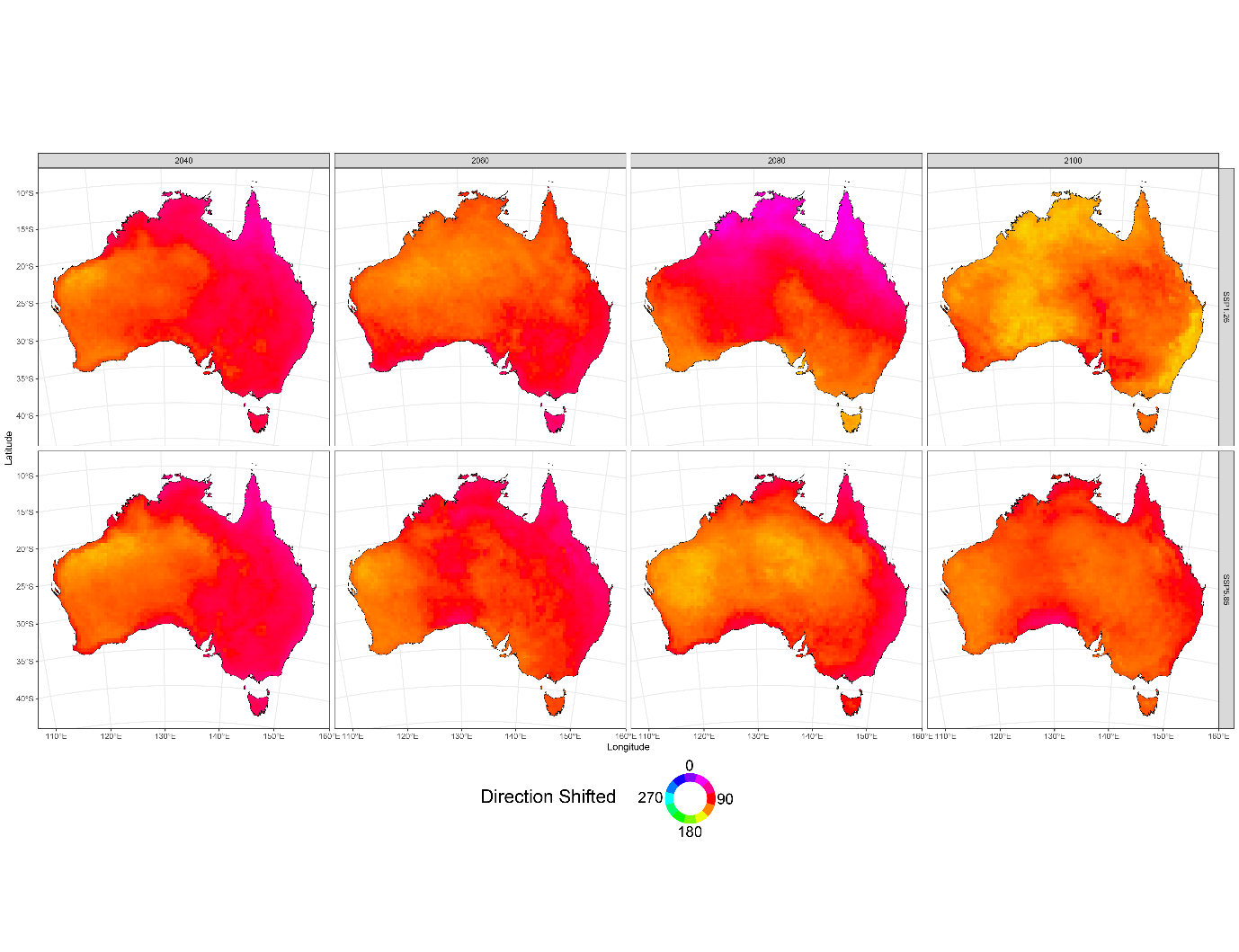

**Figure S5. Directional range shifts**. Maps showing the average direction of range centroid shifts for species per cell under different scenarios of climate change and dispersal. The colour-wheel is coded to show the angle of average range shift, with 0° representing due north: *e.g.*, redder colours indicate a shift to the east.

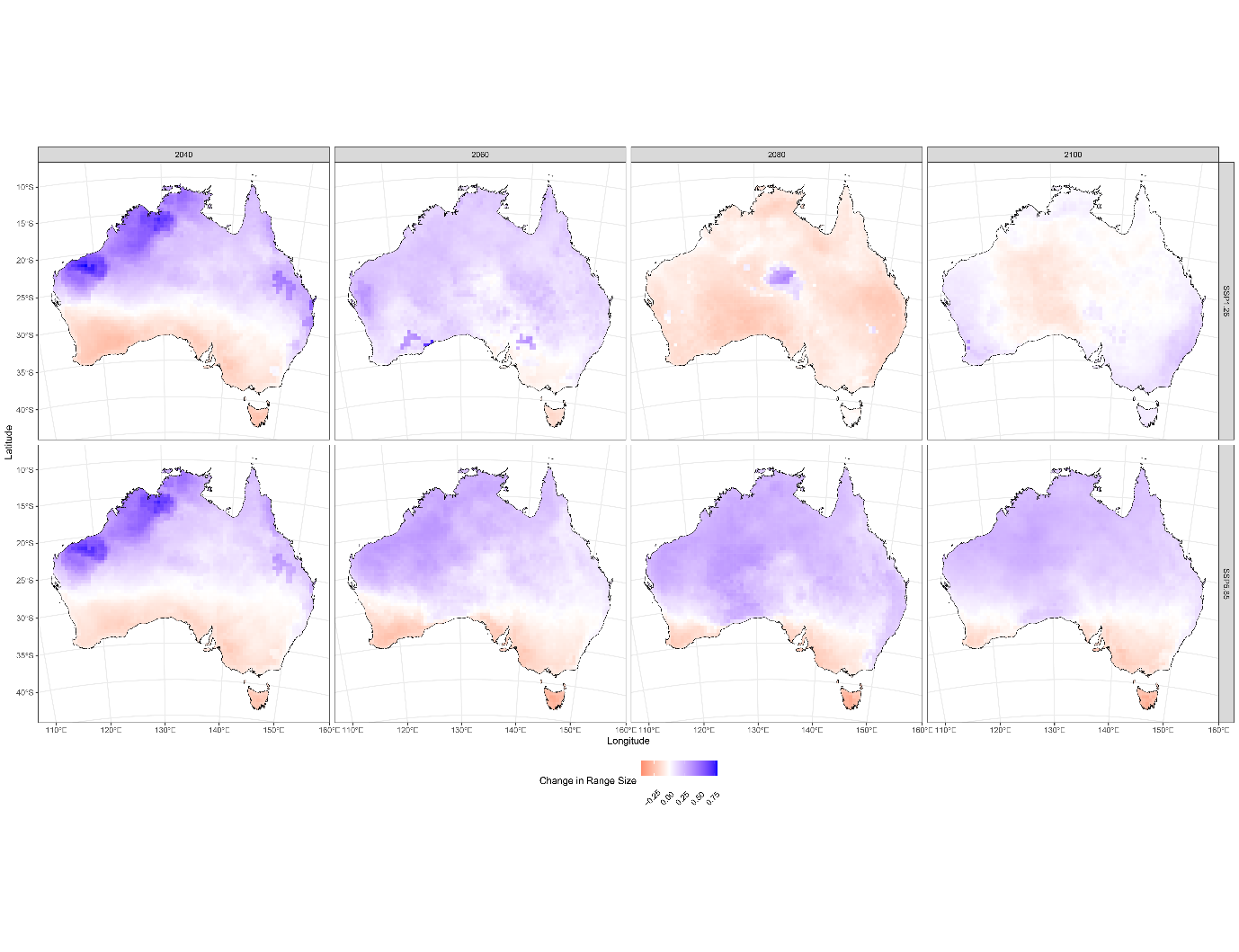

**Figure S6. Range expansions and contractions**. Maps showing the average change in range size for species per cell under different scenarios of climate change and dispersal. Negative values (red) represent range contraction compared to the previous timestep, and positive values (blue) represent range expansion compared to the previous timestep.

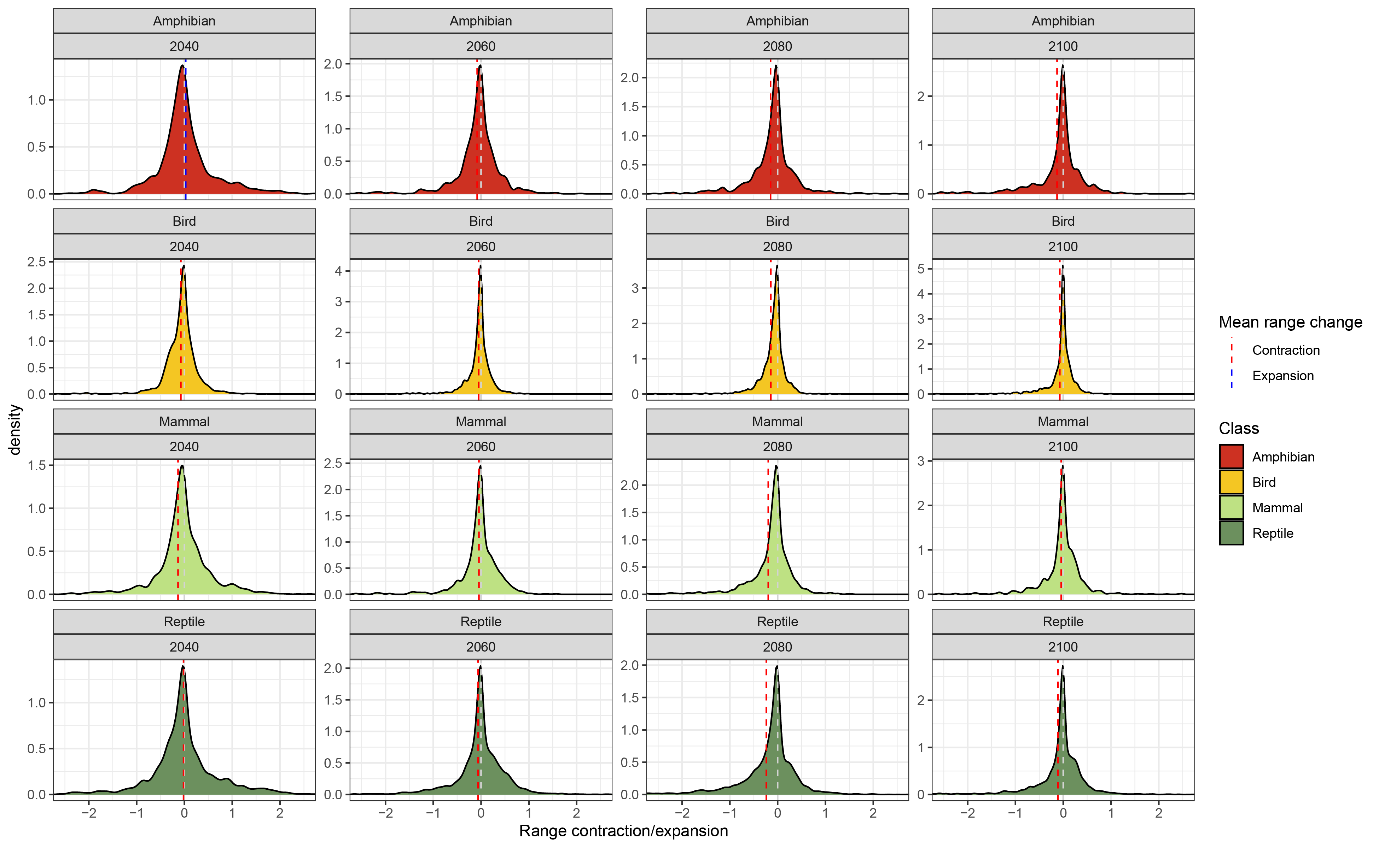

**Figure S7. Dynamics of predicted range contractions and expansions**. Density plots showing the distributions of magnitudes of range contraction/expansion in each 20-year timestep. Negative values represent range contraction compared to the previous timestep, and positive values represent range expansion compared to the previous timestep. Dashed grey lines represent 0, or no change. Dashed blue and red vertical lines represent the mean value in each timestep, blue if on average species experienced range expansions, or red if on average species experienced range contractions. Plots are truncated at -2.5 and 2.5 to improve clarity of results.

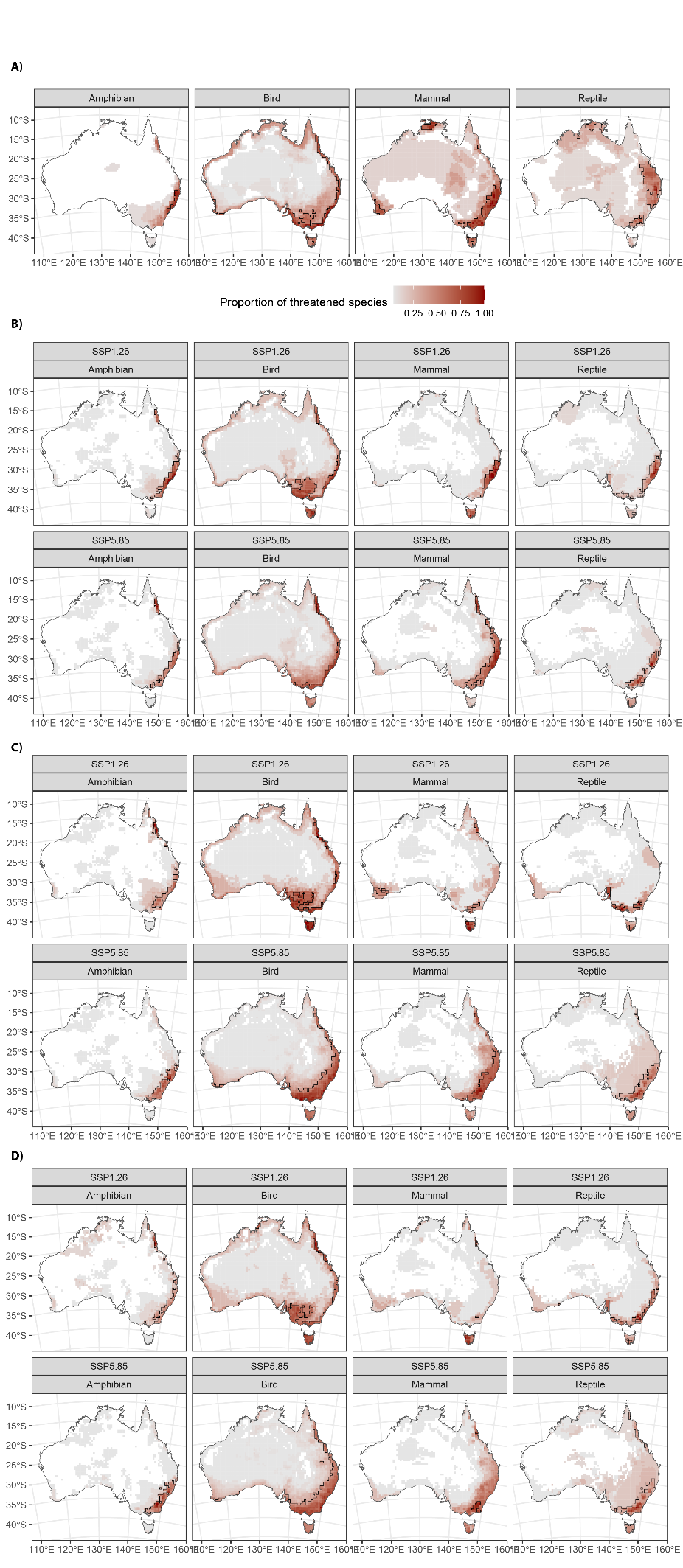

**Figure S8. Threatened species hotspots.** Threatened richness maps, showing the proportion of threatened species per cell for each tetrapod class. Thick black lines denote the hotspot (top 10^th^ percentile) for each class. A) Current threatened species richness and hotspots based on automated assessment of 1.914 species. B) Predicted threatened species richness and hotspots in 2100 under two different Shared Socioeconomic Pathways and the no dispersal scenario. C) Predicted threatened species richness and hotspots in 2100 under two different Shared Socioeconomic Pathways and the limited dispersal scenario. D) Predicted threatened species richness and hotspots in 2100 under two different Shared Socioeconomic Pathways and the limitless dispersal scenario.

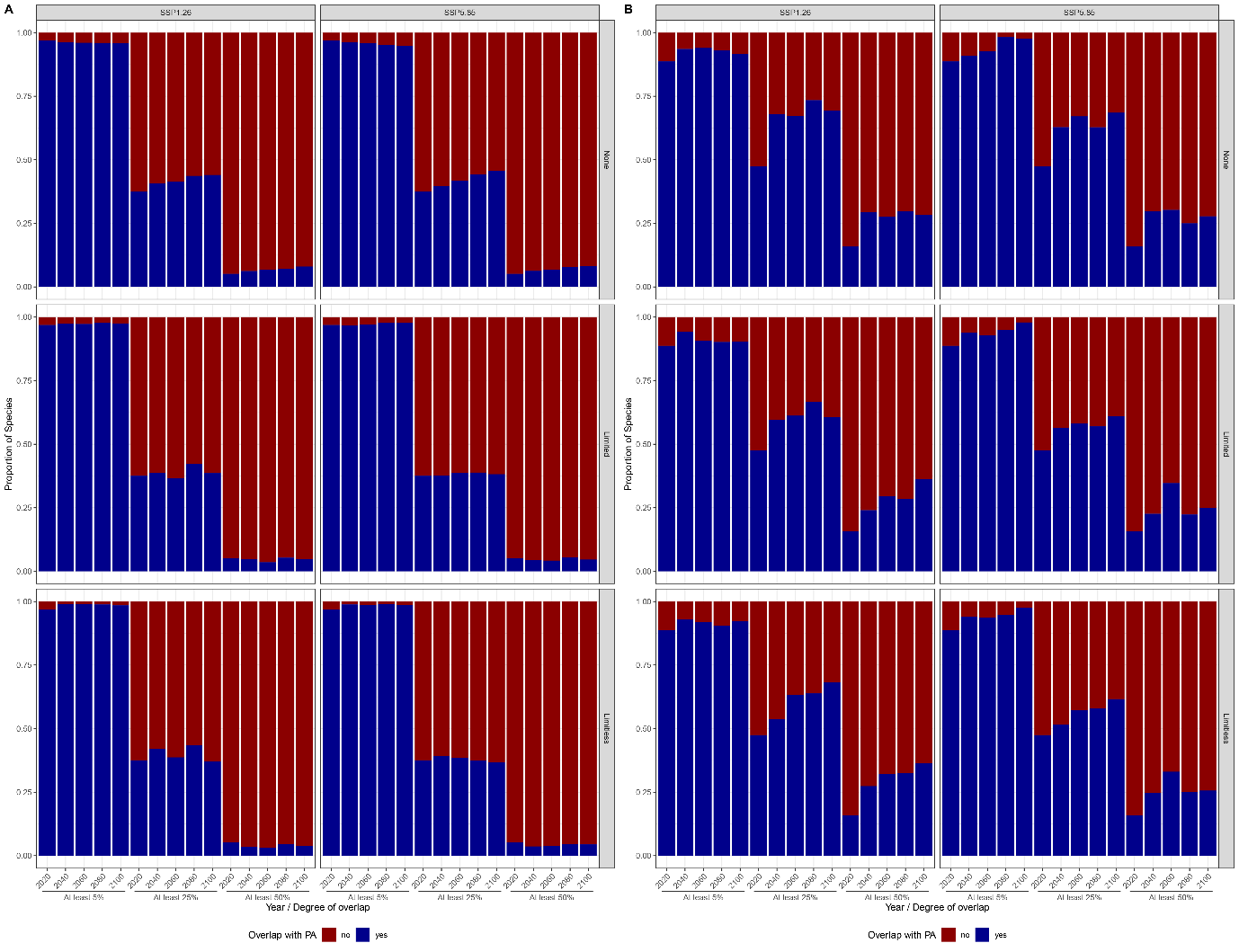

**Figure S9. Changes in levels of overlap with protected areas.** Filled barplots showing the proportion of species achieving different levels of overlap with protected areas (at least 5% of range overlap, at least 25% overlap or at least 50% overlap). Panels A and B represent the proportions of species in each level of overlap for non-threatened and threatened species, respectively.

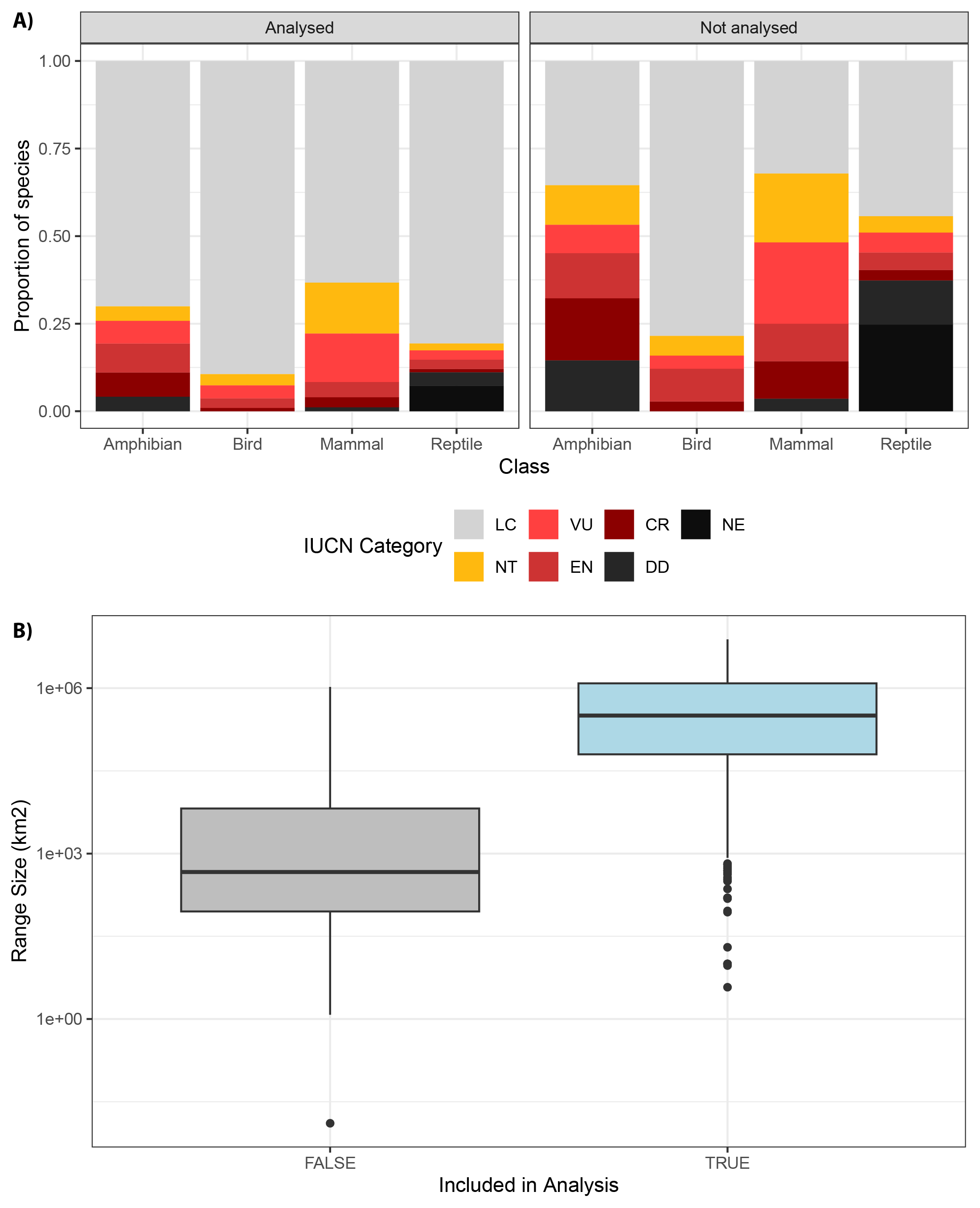

**Figure S10. Comparison of analysed species with species omitted from automated assessment**. Species were omitted from the automated assessment process either due to insufficient sample size to generate SDMs, failure to generate spatial blocks or fit SDMs, or large incongruency between SDM predicted ranges and range shapefiles. A) Bar plots comparing distributions of IUCN threat categories between analysed and non-analysed species. B) Box plot comparing the distribution of range size (in km^2^; log_10_-transformed) between analysed and non-analysed species, as calculated from range shapefiles.

**Table S1.** Mean accuracy of the XGBoost model in each stage of the automated assessment algorithm.

| **Stage** | **Accuracy** |
| --- | --- |
| Threatened vs. Non-threatened | 0.914 |
| EN/CR vs. VU | 0.604 |
| NT vs. LC | 0.937 |

**Table S2.** Summary of the features included in each stage of the XGBoost automated assessment algorithm, with a column listing the gain value associated with each feature. HPD and HML refer to human population density and human-modified land, respectively. Mean of those values refers to the average across species’ ranges, and % refers to the proportion of species’ ranges overlapping areas with human population densities ≥ 100/km^2^ and areas that are ≥ 50% human modified, respectively. Threatened categories: Vulnerable (VU), Endangered (EN), Critically Endangered (CR). Non-threatened categories: Least Concern (LC), Non-threatened (NT).

| **Feature** | **Type** | **Stage** | **Gain** |
| --- | --- | --- | --- |
| Range Size | Species traits | Threatened vs Non-threatened | 0.162 |
|  |  | NT vs LC | 0.164 |
|  |  | EN/CR vs VU | 0.226 |
| Mass |  | Threatened vs Non-threatened | 0.142 |
|  |  | NT vs LC | 0.141 |
|  |  | EN/CR vs VU | 0.116 |
| Mean human population density | Habitat | Threatened vs Non-threatened | 0.096 |
|  |  | NT vs LC | 0.070 |
|  |  | EN/CR vs VU | 0.078 |
| Mean human-modified land |  | Threatened vs Non-threatened | 0.099 |
|  |  | NT vs LC | 0.060 |
|  |  | EN/CR vs VU | 0.080 |
| % human population density |  | Threatened vs Non-threatened | 0.058 |
|  |  | NT vs LC | 0.063 |
|  |  | EN/CR vs VU | 0.035 |
| % human-modified land |  | Threatened vs Non-threatened | 0.034 |
|  |  | NT vs LC | 0.047 |
|  |  | EN/CR vs VU | 0.064 |
| Camels | Invasive species | Threatened vs Non-threatened | 0.040 |
|  |  | NT vs LC | 0.040 |
|  |  | EN/CR vs VU | 0.025 |
| Cane toads |  | Threatened vs Non-threatened | 0.066 |
|  |  | NT vs LC | 0.088 |
|  |  | EN/CR vs VU | 0.110 |
| Cats |  | Threatened vs Non-threatened | 0.087 |
|  |  | NT vs LC | 0.084 |
|  |  | EN/CR vs VU | 0.127 |
| European rabbits |  | Threatened vs Non-threatened | 0.090 |
|  |  | NT vs LC | 0.055 |
|  |  | EN/CR vs VU | 0.070 |
| Red foxes |  | Threatened vs Non-threatened | 0.057 |
|  |  | NT vs LC | 0.056 |
|  |  | EN/CR vs VU | 0.042 |
| Amphibians | Taxonomy | Threatened vs Non-threatened | 0.007 |
|  |  | EN/CR vs VU | 0.002 |
| Birds |  | Threatened vs Non-threatened | 0.003 |
|  |  | NT vs LC | 0.001 |
|  |  | EN/CR vs VU | 0.009 |
| Mammals |  | Threatened vs Non-threatened | 0.013 |
|  |  | NT vs LC | 0.097 |
|  |  | EN/CR vs VU | 0.013 |
| Reptiles |  | Threatened vs Non-threatened | 0.027 |
|  |  | NT vs LC | 0.022 |
| Dry biomes | Geography | Threatened vs Non-threatened | 0.006 |
| Temperate biomes |  | Threatened vs Non-threatened | 0.013 |
|  |  | NT vs LC | 0.013 |
|  |  | EN/CR vs VU | 0.001 |
| Tropical biomes |  | Threatened vs Non-threatened | 0.001 |

**Table S3.** Summary of post-hoc Tukey test comparing variable importance between different types of features included in the XGBoost automated assessment algorithm. The Comparison column lists the pairwise comparison (Species traits: Range Size, Mass; Habitat: Mean HPD, Mean HML, % HPD, $ HML; Introduced species: Cats, red foxes, European rabbits, cane toads, dromedary camels; Taxonomy: reptiles, mammals, birds, amphibians; Geography: dry biomes, tropical biomes, temperate biomes), the Diff column lists the difference in mean Gain between the two types of features (95% CI in parentheses), and the p column lists the adjusted *p-*value. Statistically significant comparisons are marked with an asterisk.

| **Comparison** | **Diff** | **p** |
| --- | --- | --- |
| Habitat-Geography | 0.058 (0.023, 0.093) | 0.000* |
| Introduced species-Geography | 0.065 (0.030, 0.099) | 0.000* |
| Species traits-Geography | 0.147 (0.107, 0.187) | 0.000* |
| Taxonomy-Geography | 0.012 (-0.023, 0.048) | 0.865 |
| Introduced species-Habitat | 0.007 (-0.019, 0.032) | 0.943 |
| Species traits-Habitat | 0.089 (0.056, 0.122) | 0.000* |
| Taxonomy-Habitat | -0.046 (-0.073, -0.018) | 0.000* |
| Species traits-Introduced species | 0.083 (0.051, 0.114) | 0.000* |
| Taxonomy-Introduced species | -0.052 (-0.079, -0.026) | 0.000* |
| Taxonomy-Species traits | -0.135 (-0.168, -0.101) | 0.000* |

**Table S4.** Percentages of species classified in IUCN Red List categories under future climate projections. LC: Least Concern; NT: Near Threatened; VU: Vulnerable; EN: Endangered; CR: Critically Endangered; EX: Extinct. Classifications are made under three dispersal scenarios (none, limited and unlimited), two emissions scenarios (SSP1.26, SSP5.85), and four time periods (2020-2040, 2040-2060, 2060-2080, 2080-2100).

| **Dispersal scenario** | **IUCN Red List** | **2040** | **2060** | **2080** | **2100** |
| --- | --- | --- | --- | --- | --- |
| **None** | **SSP1.26** | | | | |
|  | LC | 92.32% | 91.33% | 90.18% | 89.71% |
|  | NT | 1.57% | 1.62% | 1.36% | 1.57% |
|  | VU | 2.56% | 2.87% | 3.24% | 3.40% |
|  | EN/CR | 3.13% | 3.29% | 3.55% | 3.50% |
|  | EX | 0.42% | 0.89% | 1.67% | 1.83% |
|  | **SSP5.85** | | | | |
|  | LC | 92.11% | 89.60% | 86.52% | 84.33% |
|  | NT | 1.46% | 1.15% | 1.46% | 1.36% |
|  | VU | 2.93% | 3.40% | 3.81% | 4.34% |
|  | EN/CR | 3.45% | 4.70% | 5.33% | 4.28% |
|  | EX | 0.05% | 1.15% | 2.87% | 5.69% |
| **Limited** | **SSP1.26** | | | | |
|  | LC | 93.16% | 93.16% | 92.74% | 92.79% |
|  | NT | 1.04% | 1.15% | 0.73% | 0.99% |
|  | VU | 2.87% | 2.40% | 1.93% | 2.35% |
|  | EN/CR | 2.56% | 2.77% | 3.50% | 2.61% |
|  | EX | 0.37% | 0.52% | 1.10% | 1.25% |
|  | **SSP5.85** | | | | |
|  | LC | 92.84% | 91.48% | 89.6% | 85.79% |
|  | NT | 1.10% | 0.89% | 0.94% | 1.04% |
|  | VU | 2.98% | 2.77% | 2.56% | 3.40% |
|  | EN/CR | 3.03% | 3.81% | 3.87% | 4.28% |
|  | EX | 0.05% | 1.04% | 2.77% | 5.49% |
| **Limitless** | **SSP1.26** | | | | |
|  | LC | 94.62% | 94.36% | 93.94% | 93.68% |
|  | NT | 0.63% | 0.84% | 0.94% | 0.89% |
|  | VU | 2.19% | 1.99% | 1.52% | 2.25% |
|  | EN/CR | 2.30% | 2.61% | 2.93% | 2.56% |
|  | EX | 0.26% | 0.21% | 0.68% | 0.63% |
|  | **SSP5.85** | | | | |
|  | LC | 93.83% | 92.22% | 90.96% | 87.36% |
|  | NT | 0.84% | 0.99% | 0.78% | 1.10% |
|  | VU | 2.66% | 2.77% | 2.51% | 3.19% |
|  | EN/CR | 2.61% | 3.19% | 3.45% | 3.45% |
|  | EX | 0.05% | 0.84% | 2.30% | 4.91% |

**Table S5.** Summary of ordered logistic model predicting automatically assigned IUCN threat status as a function of threat status in the previous timestep, modelled scenario (unique combination of SSP and dispersal scenario) and taxonomic Class. Under each predictor variable are shown the factor levels, with log odds ratios (95% CI in squared parentheses) and *p*-values comparing them to the baseline levels (first row for each predictor.

| **Predictor** | **Level** | **log(OR)** | ***p*-value** |
| --- | --- | --- | --- |
| IUCN threat status in previous timestep | LC | — | — |
|  | NT | 3.7 [3.5, 3.9] | 0.000 |
|  | VU | 2.9 [2.8, 3.0] | 0.000 |
|  | EN/CR | 3.8 [3.7, 3.9] | 0.000 |
| Modelled scenario | SSP1.26 Limitless dispersal | — | — |
|  | SSP1.26 Limited dispersal | 0.03 [-0.12, 0.19] | 0.685 |
|  | SSP1.26 No dispersal | 0.80 [0.65, 0.94] | 0.000 |
|  | SSP5.85 Limitless dispersal | 0.20 [0.05, 0.35] | 0.009 |
|  | SSP5.85 Limited dispersal | 0.21 [0.06, 0.36] | 0.007 |
|  | SSP5.85 No dispersal | 1.1 [0.93, 1.2] | 0.000 |
| Class | Amphibian | — | — |
|  | Bird | -0.58 [-0.70, -0.46] | 0.000 |
|  | Mammal | 0.25 [0.13, 0.38] | 0.000 |
|  | Reptile | -0.87 [-0.99, -0.75] | 0.000 |

**Table S6.** Summary of slopes for different pairwise taxon comparisons (*e.g.*, mammals-birds) of the regression between spatially corrected Pearson’s correlation coefficients and year, from an ANCOVA with an interaction term and year and correlation coefficient scaled. The Comparison column lists the pairwise comparison, the Slope column lists the slope of the regression with year, and the p column lists the adjusted *p-*value. Statistically significant comparisons are marked with an asterisk.

| **Comparison** | **Slope** | **p** |
| --- | --- | --- |
| Amphibians-Birds | -0.006 | 0.233 |
| Amphibians-Mammals | -0.011 | 0.048* |
| Amphibians-Reptiles | 0.000 | 0.935 |
| Birds-Mammals | 0.011 | 0.038* |
| Birds-Reptiles | 0.020 | 0.000* |
| Mammals-Reptiles | 0.010 | 0.059 |

**Table S7.** Predicted species extinctions under different Shared Socioeconomic Pathways (SSPs) and dispersal scenarios. For each species, the following information is provided: common name, higher taxonomy, current IUCN assessment (if the species has a national listing under the Environment Protection and Biodiversity Conservation Act 1999 [EPBC] that is the one listed, with the global IUCN assessment in parentheses), distribution, and the year predicted to go extinct.

TABLE ATTACHED AS EXCEL FILE

**Table S8.** Sample sizes in different stages of analysis, with percentages of the total in parentheses.

| **Stage** | **Amphibians** | **Birds** | **Mammals** | **Reptiles** | **Total** | **Notes** |
| --- | --- | --- | --- | --- | --- | --- |
| Range shapefiles | 219 | 630 | 276 | 1031 | 2156 | All species known in Australia |
| Occurrence records | 217 (99%) | 622 (99%) | 275 (~100%) | 1018 (99%) | 2132 (99%) | Omitting species failing the filtering process for ALA records |
| SDM (pre) | 194 (89%) | 607 (96%) | 266 (96%) | 861 (84%) | 1928 (89%) | Omitting species with insufficient sample sizes (< 10) |
| SDM (post) | 192 (88%) | 606 (96%) | 265 (96%) | 851 (83%) | 1914 (89%) | Omitting species with failed SDMs |
| Automated Assessment (training) | 157 (72%) | 523 (83%) | 220 (79%) | 731 (71%) | 1631 (76%) | Omitting species with SDM predicted ranges ≥ 3 times range shapefiles and species with NE or DD status |

**Table S9.** Summary of ensemble SDMs, showing for each species the sample size (number of presence records used to train models), the spatial block size and K used for spatial cross-validation, three different evaluation metrics (Area Under the Receiver-Operator Curve [AUC], the continuous Boyce index, and the True Skill Statistic [TSS]) and the threshold value used for binary conversion of continuous habitat suitability projections.

TABLE ATTACHED AS EXCEL FILE
